# Multiplex, multimodal mapping of variant effects in secreted proteins

**DOI:** 10.1101/2024.04.01.587474

**Authors:** Nicholas A. Popp, Rachel L. Powell, Melinda K. Wheelock, Kristen J. Holmes, Brendan D. Zapp, Kathryn M. Sheldon, Shelley N. Fletcher, Xiaoping Wu, Shawn Fayer, Alan F. Rubin, Kerry W. Lannert, Alexis T. Chang, John P. Sheehan, Jill M. Johnsen, Douglas M. Fowler

## Abstract

Despite widespread advances in DNA sequencing, the functional consequences of most genetic variants remain poorly understood. Multiplexed Assays of Variant Effect (MAVEs) can measure the function of variants at scale, and are beginning to address this problem. However, MAVEs cannot readily be applied to the ∼10% of human genes encoding secreted proteins. We developed a flexible, scalable human cell surface display method, Multiplexed Surface Tethering of Extracellular Proteins (MultiSTEP), to measure secreted protein variant effects. We used MultiSTEP to study the consequences of missense variation in coagulation factor IX (FIX), a serine protease where genetic variation can cause hemophilia B. We combined MultiSTEP with a panel of antibodies to detect FIX secretion and post-translational modification, measuring a total of 44,816 effects for 436 synonymous variants and 8,528 of the 8,759 possible missense variants. 49.6% of possible *F9* missense variants impacted secretion, post-translational modification, or both. We also identified functional constraints on secretion within the signal peptide and for nearly all variants that caused gain or loss of cysteine. Secretion scores correlated strongly with FIX levels in hemophilia B and revealed that loss of secretion variants are particularly likely to cause severe disease. Integration of the secretion and post-translational modification scores enabled reclassification of 63.1% of *F9* variants of uncertain significance in the *My Life, Our Future* hemophilia genotyping project. Lastly, we showed that MultiSTEP can be applied to a wide variety of secreted proteins. Thus, MultiSTEP is a multiplexed, multimodal, and generalizable method for systematically assessing variant effects in secreted proteins at scale.

## Introduction

Genome sequencing and clinical genetic testing have revealed a rapidly expanding universe of human genetic variants^1–3^. However, the consequences of the vast majority of these variants with regard to human disease is unknown. Variants with insufficient evidence to interpret pathogenicity are termed Variants of Uncertain Significance (VUS)^4,5^. One way to improve interpretation of VUS is to use functional evidence derived from Multiplexed Assays of Variant Effect (MAVEs)^4,6–8^. MAVEs combine a large library of genetic variants with selection for function and high-throughput DNA sequencing to characterize the functional effects of tens of thousands of variants simultaneously^9^. Because sequencing is used as a readout, every variant protein must remain linked with its cognate variant DNA sequence. Existing yeast, phage and molecular display methods satisfy this requirement by physically linking a target protein to its cognate DNA, but generally do not work for human secreted proteins which require extensive post-translational processing for function. Secreted proteins make up approximately 10% of human genes and the number of VUS in genes encoding secreted proteins is rapidly growing, so a platform for applying MAVEs to human secreted proteins is urgently needed^10^ (**Fig. 1a,b**).

**Figure 1:**
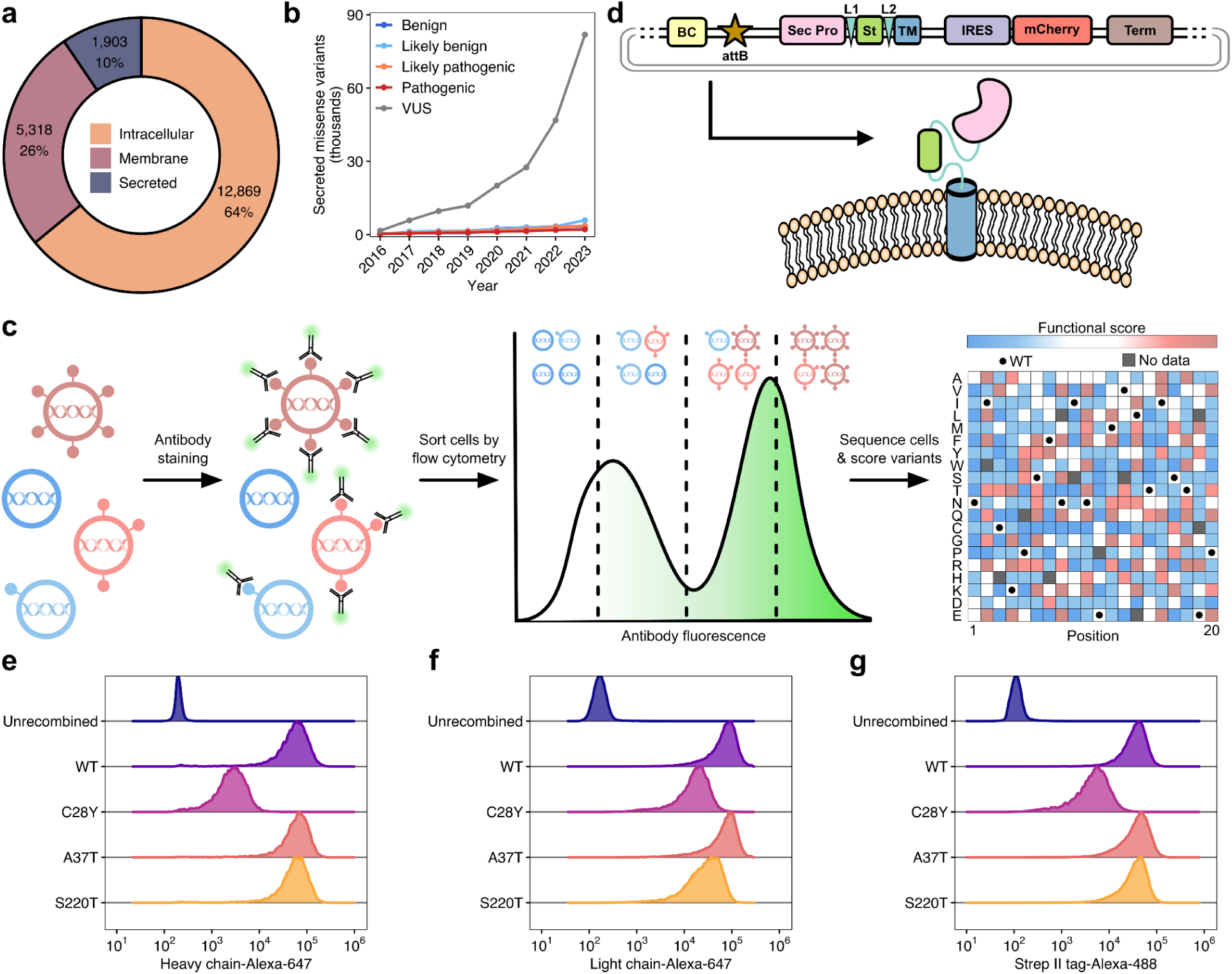
MultiSTEP enables at-scale measurement of variant effects in secreted proteins. **a.** Secreted proteins (purple) make up approximately 10% of the human proteome^10^. **b.** Missense variants in secreted proteins found in ClinVar from 2016 to 2023 colored by clinical significance. **c.** MultiSTEP retains secreted proteins on the cell surface, establishing a physical link between genotype and phenotype (left panel). Cells expressing a library of variants of the target protein are sorted into bins based upon intensity of fluorescent antibody binding, followed by deep sequencing to derive a functional score for each individual variant (middle panels). The result is a variant effect map (right panel). **d.** MultiSTEP design. Secreted protein coding sequences (pink) are cloned into an attB-containing landing pad donor plasmid. Secreted proteins are engineered to have C-terminally fused (GGGGS)_2_ flexible linkers (L1 and L2, teal) attached to a single pass transmembrane domain (TMD, blue). In between the linkers is a strep II epitope tag for surface detection (green). The construct contains an IRES (purple) driving co-transcription of an mCherry fluorophore (red) that serves as a transcriptional control. **e-g.** Flow cytometry of known well-secreted (p.A37T, p.S220T, WT) and poorly-secreted (p.C28Y) FIX variants displayed using MultiSTEP (n ∼30,000 cells per variant). Unrecombined cells do not display FIX and serve as a negative control. Fluorescent signal was generated by staining the library with either a mouse monoclonal anti-FIX heavy chain antibody (**e**), a mouse monoclonal anti-FIX light chain antibody (**f**), or a mouse monoclonal anti-strep II tag antibody (**g**), followed by staining with an Alexa Fluor-647-labeled goat anti-mouse secondary antibody.

One important secreted protein is coagulation factor IX (FIX), encoded by the *F9* gene. Normal FIX secretion and post-translational modification, including γ-carboxylation, are necessary to stop bleeding and achieve hemostasis^11–15^. Genetic variation leading to loss of *F9* function causes the bleeding disorder hemophilia B^16–21^. Undetectable FIX activity, defined as <1% (or <0.01 IU/dL), characterizes severe hemophilia B and poses high risk of developing neutralizing anti-FIX antibodies^22,23^. *F9* clinical genotyping is recommended for all individuals who have hemophilia B or are at risk to inherit a hemophilia B-causing variant^18,20,21,24,25^. Some pathogenic FIX variants are known to impact secretion (e.g. p.G253Q), while others impact FIX γ-carboxylation (e.g. p.A37D) or activity^23,26–29^. However, the effect of the vast majority of the 8,759 possible FIX missense variants is unknown. This knowledge gap is evident in the *My Life, Our Future* (MLOF) U.S. national hemophilia clinical genotyping program, where nearly half of missense variants suspected to cause hemophilia B remain classified as VUS^20,30^.

To expand MAVEs to human secreted proteins and to investigate the impact of variants on FIX secretion and post-translational modification, we developed Multiplexed Surface Tethering of Extracellular Proteins (MultiSTEP). MultiSTEP combines multiplexed human cell surface display with a genomically-encoded landing pad to measure the effects of secreted protein variants at scale^31^. We used MultiSTEP to measure both secretion and post-translational γ-carboxylation of nearly all 8,759 possible FIX missense variants using a panel of antibodies. The resulting 44,816 variant effect measurements revealed that 49.6% (n = 4,234/8,528) of FIX missense variants in our library caused loss of FIX function, with 39.5% impacting secretion, and 10.1% impacting Gla-domain γ-carboxylation or conformation. Unlike for intracellular proteins, nearly all FIX missense variants resulting in loss or gain of a cysteine diminished FIX secretion, making cysteines the most damaging substitution overall. Secretion scores correlated strongly with FIX levels in hemophilia B and revealed that loss of secretion variants are particularly likely to cause severe disease. Model-based integration of the secretion and post-translational modification scores enabled reclassification of 63.1% of *F9* VUS in the MLOF hemophilia genotyping project. Lastly, we demonstrated that MultiSTEP can be applied to multiple other clinically important secreted proteins including coagulation factors VII, VIII, and X; alpha-1-antitrypsin; plasma protease C1 inhibitor; and insulin.

## Results

### A multiplexed human cell display system compatible with secreted proteins

MultiSTEP is a human cell protein display method designed to simultaneously measure the effects of a large number of variants in secreted proteins (**Fig. 1c**). MultiSTEP combines 293-F suspension human cells with a genomically-integrated, recombinase-based landing pad to drive the expression of a single variant protein per cell^31^ (**Supplementary Fig. 1)**. Unlike most other mammalian protein display systems, the target protein’s endogenous secretion signal peptide is used, ensuring proper processing and allowing characterization of native signal peptide variants (**Fig. 1d**). An mCherry fluorescent transcriptional control is expressed via an internal ribosomal entry site (IRES)^31–33^. The C-terminus of the secreted protein is fused to a CD28 type I single-pass transmembrane domain via a multipartite, flexible linker containing a strep II tag to ensure that the protein of interest is oriented towards the outer surface of the cell and secreted using its native N-terminal signal peptide. We chose the CD28 transmembrane domain because it has been used for cell surface expression of chimeric antigen T cell receptors, is compatible with the strep II tag, and is thought to be inert^34–37^.

Using MultiSTEP, we expressed wild type (WT) FIX and detected cell surface-displayed FIX using antibodies directed against the strep II tag in the C-terminal linker, the FIX heavy chain, or the FIX light chain (**Fig. 1e-g**). Cell surface expression of p.C28Y, which reduces signal peptide cleavage efficiency and secretion, was profoundly decreased^38^ (**Fig. 1e-g**, **Supplementary Fig. 2a**). We also measured the secretion of p.C28Y using a conventional cDNA without the C-terminal linker and transmembrane domain and confirmed that it was poorly secreted in the supernatant (**Supplementary Fig. 2b**). p.A37T and p.S220T, neither of which impact FIX secretion, were present on the cell surface at levels comparable to WT ^3,16,28,39^ (**Fig. 1e-g**). Thus, MultiSTEP enabled FIX display and could be used to distinguish the effect of variants on secretion.

We measured the secretion of 8,528 of the 8,759 possible missense *F9* variants using these three secretion antibodies (**Supplementary Table 1**). We computed secretion scores for each antibody by averaging replicates (mean Pearson’s r = 0.95), normalizing such that a score of 1 indicated wild type-like secretion and a score of 0 equaled the median of the lowest-scoring 5% of missense variants (**Fig. 2a-c, Supplementary Fig. 3, Supplementary Fig. 4, Supplementary Fig. 5a**). Synonymous variants had secretion scores similar to WT, and missense variants were distributed bimodally, similar to previous multiplexed measurements of cytoplasmic protein variant abundance^33,40,41^ (**Fig. 2d,e, Supplementary Fig. 5b**). Measurement of individual variant antibody binding strongly correlated with the MultiSTEP-derived secretion scores for seven randomly selected variants spanning the secretion score range (**Fig. 2f,g, Supplementary Fig. 5c**). MultiSTEP secretion scores also correlated with the quantity of untethered FIX secreted from 293-F cells for eight variants tested (Pearson’s r = 0.93, **Supplementary Fig. 2b**). In adherent HEK-293T cells, non-enzymatic cell dissociation followed by measurement of cell surface levels of 20 surface-tethered FIX variants correlated with our MultiSTEP-derived secretion scores (Pearson’s r = 0.9, **Supplementary Fig. 2c**).

**Figure 2:**
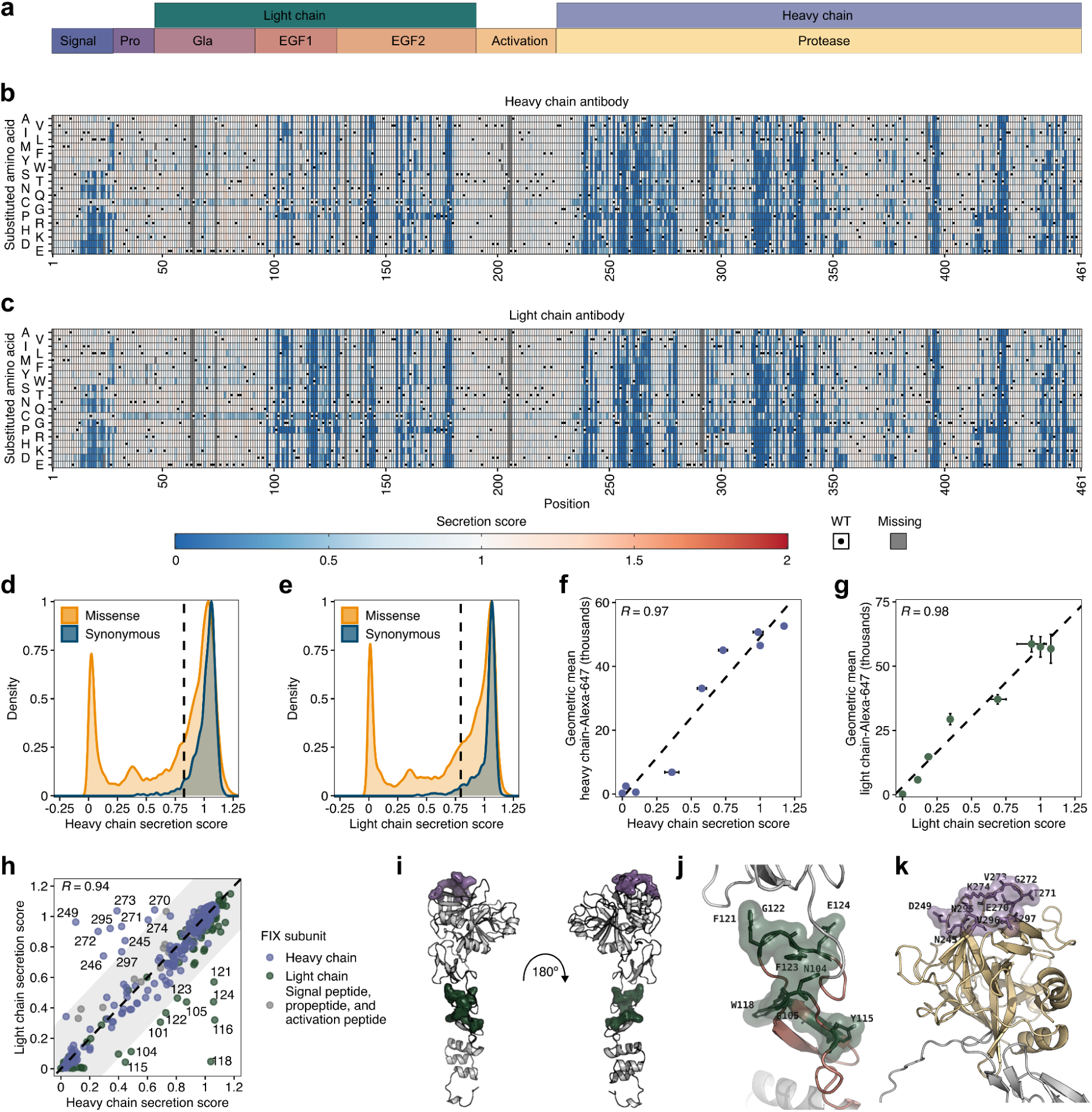
17,927 MultiSTEP-derived secretion scores for 8,964 factor IX variants. **a.** Factor IX domain and chain architecture. Signal: Signal peptide. Pro: Propeptide. Gla: Gla domain. EGF1: Epidermal growth-like factor 1 domain. EGF2: Epidermal growth-like factor 2 domain. Activation: Activation peptide. Protease: Serine protease domain. **b-c.** Heatmaps showing FIX heavy chain secretion scores (**b**) or FIX light chain secretion scores (**c**) for nearly all missense FIX variants. Heatmap color indicates antibody score from 0 (blue, lowest 5% of scores) to white (1, WT) to red (increased scores). Black dots indicate the WT amino acid. Missing data scores are colored gray. **d-e.** Density distributions of heavy chain (**d**) or light chain (**e**) secretion scores for FIX missense variants (orange) and synonymous variants (blue). Dashed line denotes the 5th percentile of the synonymous variant distribution. **f-g.** Scatter plots comparing MultiSTEP-derived heavy chain (**f**) or light chain (**g**) secretion scores for seven different FIX variants (p.C28Y, p.A37T, p.G58E, p.E67K, p.C134R, p.S220T, and p.H267L), WT, and an unrecombined negative control to the geometric mean of Alexa Fluor-647 fluorescence measured using flow cytometry on cells expressing each variant individually. The p.E67K missense variant is not present in (**g**). **h.** Scatter plot of median MultiSTEP-derived heavy chain and light chain secretion scores at each position in FIX. Points are colored by chain architecture, using the same color scheme as (**a**). Black dashed line indicates the line of perfect correlation between secretion scores. Pearson’s correlation coefficient is shown. Gray background indicates <0.3 point deviation from perfect correlation. Points with median positional scores outside gray background are labeled with their corresponding FIX position. **i.** AlphaFold2 model of mature, two-chain FIX (positions 47-191 and 227-461). Positions labeled in (**h**) are shown as colored surfaces where color corresponds to the FIX heavy chain (purple) or light chain (green) putative epitope positions. **j.** Magnified view of FIX EGF1 domain in the light chain (orange). Putative epitope positions with discordant light and heavy chain antibody scores (**h-i**) are shown as a green colored surface with visible amino acids labeled. **k.** Magnified view of FIX serine protease domain in the heavy chain (yellow). Putative epitope positions (**h-i**) are shown as a purple colored surface with visible amino acids labeled.

The heavy chain, light chain, and strep tag antibody-derived secretion scores were nearly identical at most positions, though the strep tag demonstrated a wider distribution of scores for synonymous variants (**Fig. 2h, Supplementary Fig. 5d-f**). A few positions showed antibody-specific differences: the median heavy chain antibody-derived secretion scores were lower than the median light chain secretion scores at 10 positions in the heavy chain, and the median light chain antibody-derived secretion scores were lower than median heavy chain secretion scores at 10 positions in the light chain (**Fig. 2h**). These two sets of positions occupied discrete, solvent-accessible sites on FIX structure, strongly suggesting that these positions comprise each antibody’s epitope (**Fig. 2i-k**). To prevent epitope effects from impacting subsequent analyses, we excluded secretion scores derived from each antibody at their respective epitope positions.

We surveyed all positions within the light chain and heavy chain for short-range effects on antibody binding using changepoint analysis (**Supplementary Fig. 6a-c**). For the light chain epitope, residues within 9.15 angstroms of our identified epitope showed a small (< 0.3 secretion score difference) decrease in light chain secretion scores relative to heavy chain. The same effect was found for residues within 5.71 angstroms of the heavy chain epitope.

### MultiSTEP clarifies the role of the signal peptide and cysteine effects on secretion

Signal peptides direct protein secretion and vary in length and sequence^42–44^. Three distinct functional regions are conserved amongst human signal peptides: the N-region, which is weakly positively charged; the H-region, which contains the hydrophobic helix that binds to a signal recognition particle (SRP) to initiate translocation into the ER; and the C-region, which breaks the h-region helix and contains a canonical AxA signal peptide cleavage motif^43,45–47^. Variants had distinct effects in each region of the signal peptide (**Fig. 3a**). For example, positions p.A26 and p.C28 within the AxA motif tolerated substitution with small amino acids, consistent with these positions occupying shallow hydrophobic pockets near the active site of the signal peptidase^47^. Surprisingly, hydrophobic variants in the spacer of the AxA motif caused significant loss of secretion, potentially by extending the H-region hydrophobic helix and preventing cleavage^43,48,49^.

**Figure 3:**
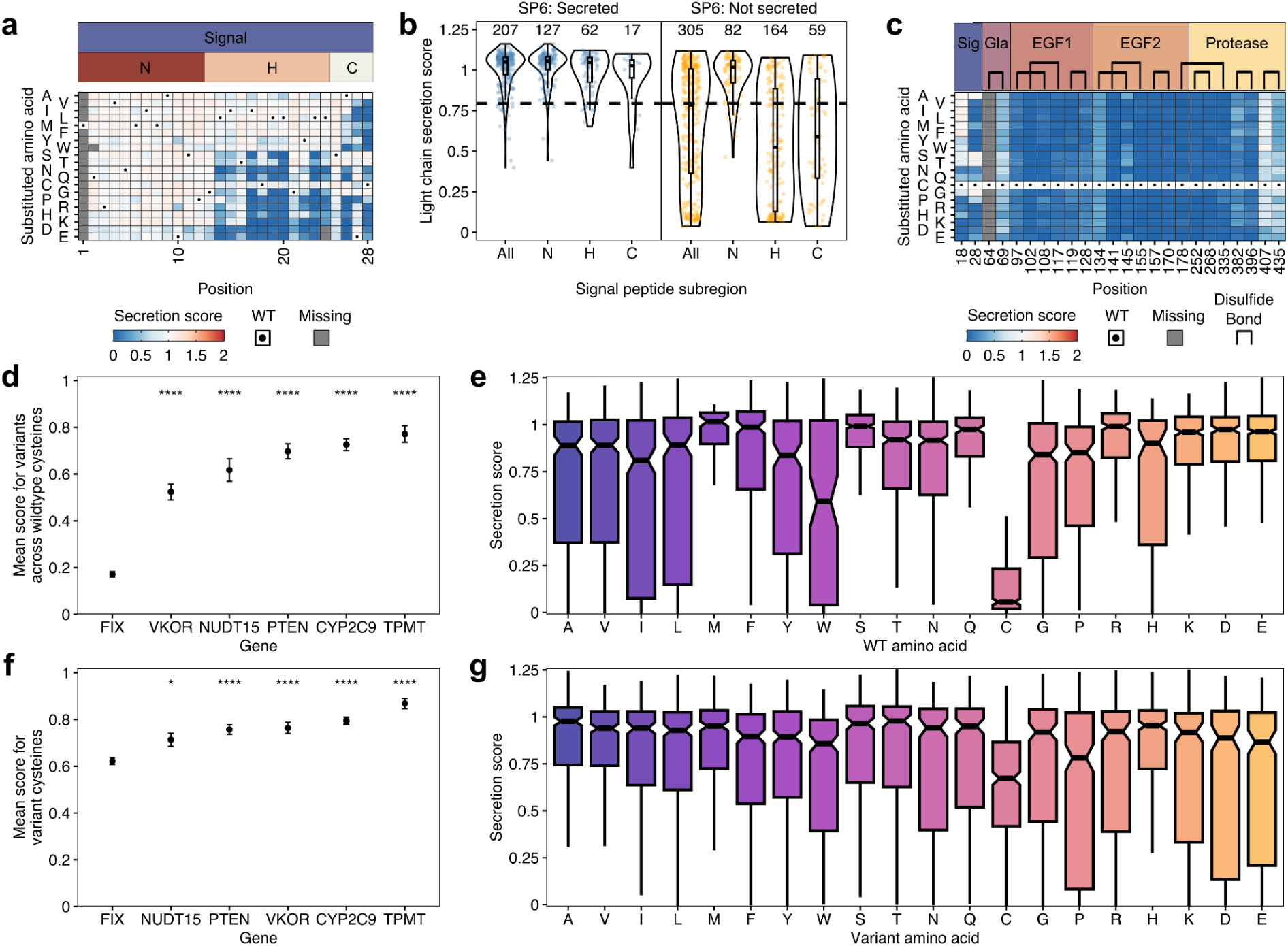
MultiSTEP reveals biochemical constraints on secretion. **a.** Predicted signal peptide regions for WT FIX from SignalP 6.0, indicated by color (top). Heatmap showing FIX heavy chain secretion scores for signal peptide variants (bottom). Heatmap color indicates antibody score from 0 (blue, lowest 5% of scores) to white (1, WT) to red (increased antibody scores). Black dots indicate the WT amino acid. Missing scores are gray. N: N-region; H: H-region; C: C-region. **b.** Secretion antibody scores for variants predicted by SignalP 6.0 to be either secreted or not secreted, throughout signal peptide and within sub-regions. Violin plot shows distribution of points with an inset box plot and horizontal lines representing the 25^th^, 50^th^, and 75^th^ percentiles. Dashed horizontal line is the 5^th^ percentile of the synonymous secretion score distribution. Number of variants in each class is labeled above the violin plot. SP6: SignalP 6.0; N: N-region; H: H-region; C: C-region. **c.** FIX cysteine variants colored by domain architecture (top). Sig: Signal peptide. Gla: Gla domain. EGF1: Epidermal growth-like factor 1 domain. EGF2: Epidermal growth-like factor 2 domain. Protease: Serine protease domain. Disulfide bridges in WT FIX are denoted by black connecting lines (p.C18 and p.C28 do not form a disulfide bridge)^14,159–161^. Heatmap of FIX heavy chain secretion scores for variants at positions with a cysteine in the WT sequence, colored as in (**b**) (bottom). **d.** Mean variant effect scores for all loss-of-cysteine substitutions for different proteins. For FIX, the average secretion score determined by MultiSTEP is shown. For all other proteins, the average abundance score measured using VAMP-seq is shown^33,40,41,58^. Points indicate mean abundance or secretion score for all scored variants at all WT cysteine positions. Error bars show standard error of the mean. Asterisks indicate level of statistical significance as determined by Bonferroni-corrected pairwise two-sided t-test to FIX variant scores. **** = *p* <0.0001. **e.** Box plots of secretion scores for all missense variants across all positions with the indicated WT amino acid. **f.** Mean variant effect scores for all gain-of-cysteine substitutions for the same set of proteins as in (**d**). Points indicate mean abundance or secretion score for all gain-of-cysteine variants. Error bars show standard error of the mean. Asterisks indicate level of statistical significance as determined by Bonferroni-corrected pairwise two-sided t-test to FIX variant scores. * = *p* <0.05, **** = *p* <0.0001. **g.** Box plots of secretion scores for all missense substitutions of the indicated variant amino acid across all positions.

93.2% (n = 193/207) of variants predicted to be secreted by the signal peptide classifier SignalP 6.0 were also secreted in our assay, with few notable exceptions^50^ (e.g. p.E27P). However, 56.4% variants predicted to not to be secreted by SignalP were secreted normally in our assay, including p.M8T, a known benign variant^20^ (n = 172/305, **Fig. 3b**). The N-region of the FIX signal peptide was largely tolerant of most substitutions aside from the introduction of negatively-charged amino acids (D, E) which led to mild loss of secretion (**Fig. 3a**). These results are consistent with our understanding of the N-region as being both positively charged and less conserved relative to the other signal peptide regions^51,52^. Indeed, nearly all (87.8%, n = 72/82) N-region variants predicted to be poorly secreted by SignalP had WT-like secretion scores. 34.2% (n = 56/164) of H-domain and 35.6% (n = 21/59) of C-domain variants were similarly misclassified by SignalP (**Fig. 3b**). Our data support that, at least for FIX, while SignalP can predict the existence of the signal peptide and cleavage site, it has poor specificity to predict variant effects on secretion.

Many secreted proteins require disulfide bonds to establish and maintain structure, and variants that remove important cysteines can cause disease^53–56^. In FIX, 22 of the 24 native cysteines are disulfide-bonded, including positions p.C178 and p.C335 which form an interdomain disulfide bond critical to maintaining activated FIX integrity after proteolytic cleavage of the activation peptide^57^. Most variants at WT cysteine positions dramatically reduced secretion, especially in the EGF1, EGF2, and serine protease domains, where most positions were completely intolerant of substitution (**Fig. 3c**). This detrimental impact of substitution on cysteines is markedly different from the much more modest effects of substitution of WT cysteines in non-secreted cytoplasmic and transmembrane proteins (**Fig. 3d, Supplementary Fig. 7a**). WT cysteine positions were overwhelmingly the most intolerant of substitution in FIX compared to other positions (**Fig. 3e**).

Missense variants that introduced new cysteines also caused loss of secretion throughout FIX (**Fig. 2b-c**). Introduction of a cysteine was the most damaging substitution, unlike cytoplasmic and membrane-associated proteins where proline or tryptophan were generally most damaging^33,40,41,58^ (**Fig. 3f-g, Supplementary Fig. 7b**). Cysteine substitutions were deleterious even within the activation peptide, a highly flexible loop removed during FIX activation where other substitutions did not appreciably impact secretion (**Fig. 2a-c**). Taken together, the deleterious effect of either removing a WT cysteine or introducing a new cysteine strongly suggests that altering the number or location of cysteines in FIX is fundamentally detrimental to FIX secretion, likely through changes in disulfide bonding patterns or protein folding differences under oxidative stress. These results contrast with cytoplasmic proteins, where removing or introducing cysteines has a more modest effect.

Evolutionary conservation also influences tolerance to substitution. While poorly conserved positions in FIX were generally tolerant of variation, variants at highly conserved positions were 5.46 times more likely to impact secretion^59^ (p < 2.2 x10-16, two-sided Fisher’s exact test, **Supplementary Fig. 8a,b**). 19 variants at poorly conserved positions impacted secretion severely (secretion score < 0.1), and 16 of these variants were found in the signal peptide, which varies widely in sequence and in length across proteins. Conversely, 87 variants at highly conserved positions had WT-like secretion scores. Many of these conserved positions are associated with specific molecular functions not captured by our secretion assay including γ-carboxylation, activation peptide cleavage, enzymatic activity, and partner binding. Thus, the effects of substitutions on FIX secretion are largely consistent with conservation.

### MultiSTEP quantifies the impact of variation on FIX γ-carboxylation and Gla domain conformation

Twelve glutamates in the FIX Gla domain are γ-carboxylated, and γ-carboxylation is required for FIX activation and function. Some of these γ-carboxylated glutamates coordinate a calcium-dependent conformational change in the Gla domain, creating a three-helix structure that exposes the ω-loop^11–14,60–67^. Only 17 of the 106 hemophilia B-associated variants in either the propeptide and Gla domain have been mechanistically shown to alter function of the FIX Gla domain^16,17,23,28,39,68–70^.

We used two different γ-carboxylation-sensitive, anti-Gla domain antibodies to interrogate the effect of FIX variants on γ-carboxylation. One is a FIX-specific antibody that recognizes the γ-carboxylation-dependent exposure of the ω-loop^64,71,72^. Thus, binding of this antibody depends both on γ-carboxylation and Gla domain conformation. The other is a Gla-motif-specific antibody that interacts with γ-carboxylated glutamates in a conserved ExxxExC motif present in the Gla domain of multiple carboxylated proteins, including FIX^28,73,74^ (**Supplementary Fig. 9a**). As expected, WT FIX-displaying cells strongly bound both antibodies. Incubation of cells with warfarin, a drug that inhibits γ-carboxylation, prior to induction of WT FIX expression eliminated binding of both γ-carboxylation-sensitive antibodies, confirming that they only detect γ-carboxylated FIX^75^ (**Supplementary Fig. 9b,c**).

We used the FIX-specific and Gla-motif γ-carboxylation-dependent antibodies to generate two different γ-carboxylation scores for nearly all possible FIX missense variants (**Fig. 4a-c**). Like for secretion, missense variant γ-carboxylation scores were bimodally distributed and synonymous variants scored similarly to WT for both antibodies (**Fig. 4d-e**). γ-carboxylation scores correlated well with secretion scores except in the propeptide and Gla domain, where some variants impacted γ-carboxylation without affecting secretion (Pearson’s r = 0.85, **Fig. 4f, Supplementary Fig. 9d-i**). Thus, in our system loss of FIX γ-carboxylation did not impact secretion (**Supplementary Fig. 9f,i**). Indeed, of the 1,154 missense variants in the propeptide and Gla domains that we assayed, 44.6% (n = 515) were associated with low carboxylation scores but normal secretion scores.

**Figure 4:**
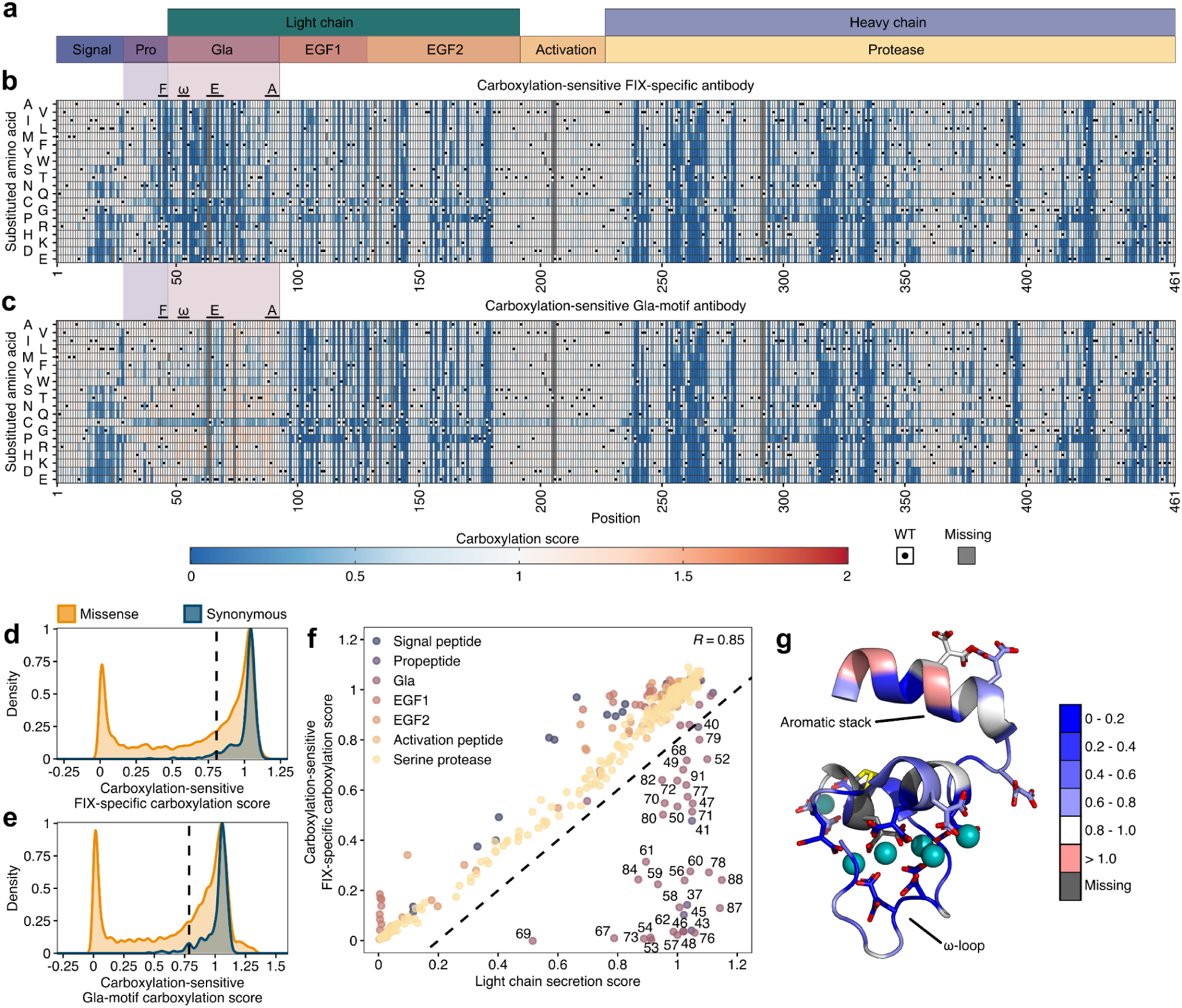
MultiSTEP enables measurement of variant effects on FIX post-translational modification. **a.** Factor IX domain and chain architecture. Signal: Signal peptide. Pro: Propeptide. Gla: Gla domain. EGF1: Epidermal growth-like factor 1 domain. EGF2: Epidermal growth-like factor 2 domain. Activation: Activation peptide. Protease: Serine protease domain. **b-c.** Heatmaps showing carboxylation-sensitive FIX-specific carboxylation scores (**b**) or carboxylation-sensitive Gla-motif carboxylation scores (**c**) for nearly all missense FIX variants. Heatmap color indicates antibody score from 0 (blue, lowest 5% of scores) to white (1, WT) to red (increased antibody scores). Black dots indicate the WT amino acid. Missing data scores are colored gray. Furin cleavage site (F), ω-loop (ω), ExxxExC motif (E), and aromatic stack (AS) are annotated above (**b**) and (**c**). For zoomed-in heatmaps on the propeptide and Gla domains of FIX, please refer to **Supplementary Fig. 9d-i**. **d-e.** Density distributions of carboxylation-sensitive FIX-specific (**d**) or carboxylation-sensitive Gla-motif (**e**) carboxylation scores for FIX missense variants (orange) and synonymous variants (blue). Dashed line denotes the 5th percentile of the synonymous variant distribution. **f.** Scatter plot of median MultiSTEP-derived carboxylation-sensitive FIX-specific carboxylation scores and light chain secretion scores at each position in FIX. Points are colored by domain architecture, using the same color scheme as **a**. Black dashed line indicates >0.2 point deviation threshold from perfect correlation between carboxylation and secretion scores. Points with deviation greater than this threshold are labeled with their corresponding FIX position. Pearson’s correlation coefficient is shown. **g.** Crystal structure of FIX Gla domain (positions 47-92)^14^. Disulfide bridges and γ-carboxylated glutamates are shown as sticks. Calcium ions are shown as teal spheres. Structure is colored according to the ratio of median positional carboxylation-sensitive FIX-specific carboxylation score to median positional FIX light chain secretion score. Missing positions are colored gray. Disulfide bridge side chains are colored yellow.

The FIX propeptide mediates binding of gamma-glutamyl carboxylase, the enzyme that γ-carboxylates the Gla domain prior to secretion^60–63^. Propeptide positions p.F31, p.A37, p.I40, p.L41, p.R43, p.K45, and p.R46 are important for Gla domain γ-carboxylation, though the importance of p.R43, p.K45, and p.R46 is contested^16,17,28,29,76–78^. Some substitutions at the highly conserved position p.A37 in the propeptide (e.g. p.A37D) prevent gamma-glutamyl carboxylase binding and thus Gla domain γ-carboxylation^28,79^. In our assay, 12 of the 19 substitutions at position p.A37, including p.A37D, reduced γ-carboxylation scores for both antibodies. Serine, threonine, and glutamate substitutions at position p.I40, which is conserved across all Gla-containing coagulation proteins, also reduced γ-carboxylation scores (**Fig. 4a-c**, **Supplementary Fig. 9d-f**). Thus, like position p.A37, position p.I40 appears to be important for γ-carboxylation. The propeptide furin cleavage motif (p.R43 to p.R46) harbors numerous severe hemophilia B-associated variants^23^ (**Fig. 4a-c, Supplementary Fig. 9d-f**). Variants in the furin cleavage motif can block propeptide cleavage, leading to its retention^80,81^. The impact of propeptide retention on Gla domain γ-carboxylation remains controversial, with some data suggesting that γ-carboxylation itself is blocked by the retained propeptide and other data suggesting that the retained propeptide alters the conformation of the ω-loop by disrupting electrostatic interactions within the Gla domain^11,28,76,78,80–82^. In our assay, variants in the furin cleavage motif had low scores for the FIX-specific γ-carboxylation antibody but WT-like scores for the Gla-motif antibody and the secretion antibodies (**Fig. 2b,c**; **Fig. 4b,c, Supplementary Fig. 9d-f**). One possible explanation for discordance between the FIX-specific and Gla-motif antibody scores at propeptide cleavage motif positions is that these substitutions perturb ω-loop conformation but permit γ-carboxylation of the glutamates in the conserved Gla-motif.

We compared the FIX-specific Gla domain and Gla-motif γ-carboxylation antibody scores for each variant within the Gla domain and found that many variants in the ω-loop (positions p.N49 to p.Q57) had WT-like scores for both antibodies (**Fig. 4b,c, Supplementary Fig. 9g-i**). The lowest FIX-specific Gla domain antibody γ-carboxylation scores resulted from substitutions at positions involved in the coordination of calcium ions (p.N48, p.E53, p.E54, p.E67, p.E73, and p.E76) in the Gla domain core^14^ (**Fig. 4g, Supplementary Fig. 9g-i**). Variants at positions p.F87 and p.W88, which form an aromatic stack required for divalent cation-induced folding of the Gla domain, had low FIX-specific γ-carboxylation scores but WT-like secretion scores likely due to disrupted γ-carboxylation, Gla domain conformation, or both^83,84^ (**Fig. 4f, Supplementary Fig. 9g-i**). Unexpectedly, variants at half of the γ-carboxylated glutamate residues in the Gla domain (p.E66, p.E72, p.E79, p.E82, p.E86) did not impact or only modestly reduced FIX-specific Gla domain and Gla-motif antibody γ-carboxylation scores (**Fig. 4b,c,g, Supplementary Fig. 9g-i**). These glutamate residues coordinate magnesium ions rather than calcium ions at physiologic concentrations^13,85–87^ (**Fig. 4g**). In contrast, many Gla domain variants outside of the ExxxExC conserved Gla motif (spanning p.E63 to p.C69) had WT-like Gla-motif antibody scores despite their low FIX-specific Gla-domain antibody scores (**Fig. 4b,c, Supplementary Fig. 9a,d-i**). Within the conserved Gla-motif, variants at positions p.R62, p.E66, p.E67, and p.C69 showed similar score reductions for both antibodies. Likewise, low variant scores were detected for both carboxylation-sensitive antibodies around the furin cleavage site in the FIX propeptide (p.R43, p.K45, and p.R46) (**Supplementary Fig. 9d-i**).

### Secretion scores reflect circulating FIX levels

We previously established that many pathogenic missense variants in intracellular proteins act by disrupting protein abundance^33,40,41,58^. However, for secreted proteins, the predominance of loss of secretion among pathogenic variants remains largely unexplored. Thus, we investigated whether MultiSTEP-derived FIX secretion scores could predict circulating FIX antigen levels in individuals with hemophilia B. Secretion scores correlated strongly with plasma FIX antigen levels reported in the European Association for Haemophilia and Allied Disorders (EAHAD) public database^23^ (n = 85, Pearson’s r = 0.76; **Fig. 5a**). Removal of variants at epitope-adjacent positions did not appreciably affect the correlation between FIX antigen and MultiSTEP secretion scores (Pearson’s r = 0.75; **Supplementary Fig. 10a**). We classified variants with secretion scores less than the 5th percentile of synonymous variants as lowly secreted^33,41,58^. 90.2% (n = 37/41) of lowly secreted variants had low circulating FIX levels, defined as <40% of pooled normal plasma^23^. The four lowly secreted variants, p.R75Q, p.R191C, p.R379G, and p.A436V, that had normal plasma FIX levels *in vivo* had secretion scores or reported FIX levels close to the assay thresholds, suggesting measurement error as the likely explanation (**Supplementary Table 2**). Conversely, five variants, p.F71S, p.D93N, p.A173V, p.Q241H, and p.N393T had markedly increased secretion scores relative to their FIX antigen levels (**Supplementary Table 2**). These variants occurred throughout the protein, outside of known binding sites for extravascular compartments or clearance binding partners^88–91^. These variants could be secreted normally but have a shortened half-life *in vivo*, which would be missed by our assay.

**Figure 5:**
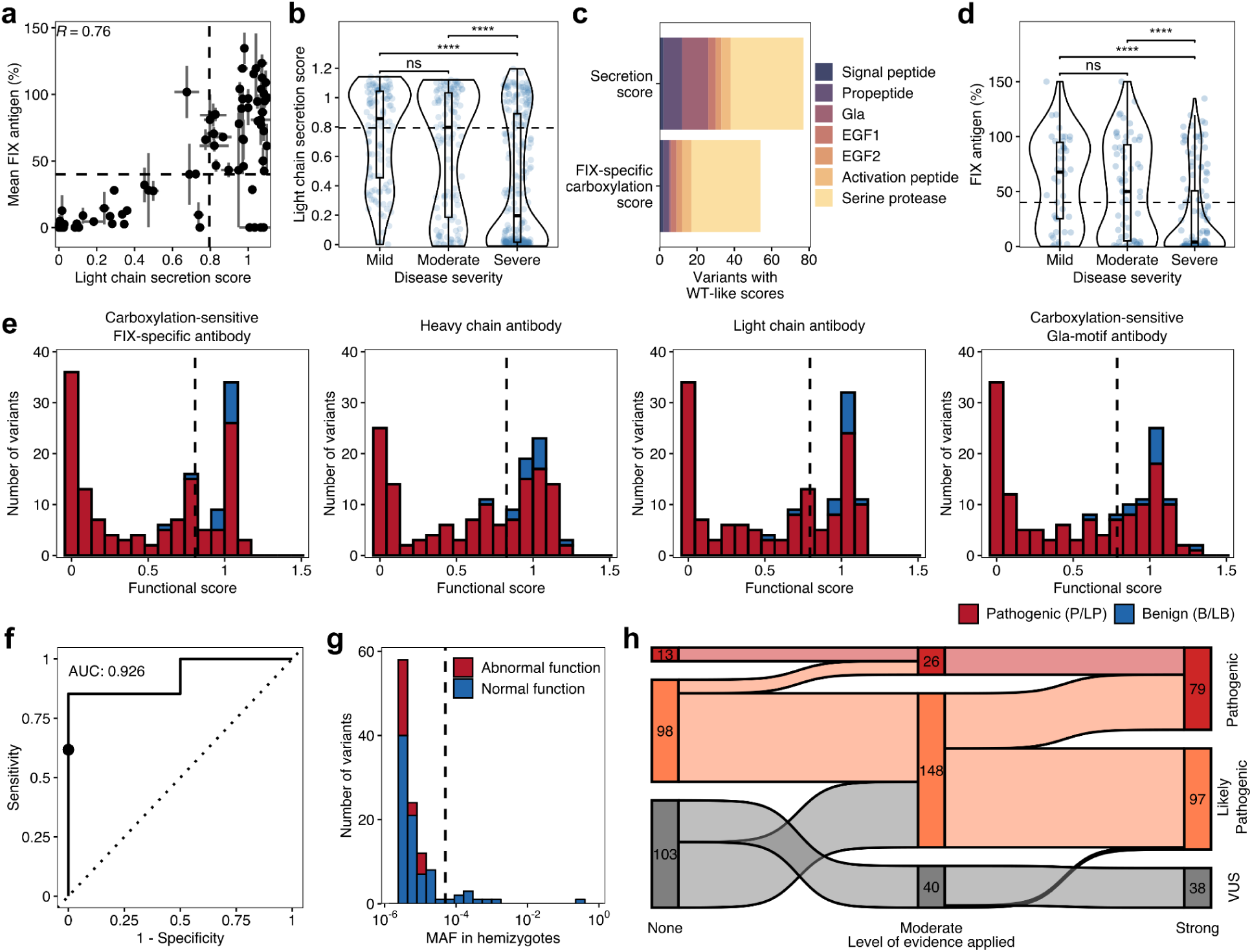
Secretion and gamma-carboxylation scores reveal clinical features of hemophilia B and enable variant reinterpretation. **a.** Scatter plot of light chain secretion scores and FIX plasma antigen from individuals with hemophilia B in the EAHAD database^23^. Horizontal solid lines indicate standard error of the mean for light chain secretion scores. Vertical solid lines indicate standard error of the mean for FIX plasma antigen levels across individuals with hemophilia B harboring the same variant. Dashed horizontal line is 40% FIX plasma antigen. Dashed vertical line is the 5^th^ percentile of the synonymous secretion score distribution. LOESS line of best fit depicted in red with 95% confidence interval shaded in gray. **b.** Comparison of EAHAD individual hemophilia B severity with light chain secretion scores from MultiSTEP. Violin plot shows distribution of points with an inset box plot representing the 25^th^, 50^th^, and 75^th^ percentiles. Dashed horizontal line is the 5^th^ percentile of the synonymous secretion score distribution. To compare median secretion scores across disease severities, a Kruskal–Wallis test adjusted for multiple comparisons by post-hoc Dunn’s test was performed. Asterisks indicate the level of statistical significance. ns = p >0.05. **** = *p* <0.0001. **c.** Severe hemophilia B disease-associated variants with WT-like light chain-derived secretion scores or FIX-specific γ-carboxylation antibody scores is shown. Bars are colored according to the number of variants located in the indicated domain. **d.** Comparison of EAHAD individual hemophilia B severity with EAHAD FIX plasma antigen levels. Dashed horizontal line is 40% FIX plasma antigen. To compare median FIX plasma antigen levels across disease severities, a Kruskal-Wallis test followed by post-hoc Bonferroni-corrected Dunn’s test was performed. Asterisks indicate the level of statistical significance. ns = p >0.05. **** = *p* <0.0001. **e.** Histograms of multiplexed functional scores for *F9* missense variants of known effect curated from ClinVar, gnomAD, and MLOF. Color indicates clinical variant interpretation. Data from four antibodies are shown. Dashed vertical line indicates the 5^th^ percentile of synonymous variants used as a threshold for abnormal function. **f.** Receiver-operator curve for random forest classifier for identifying abnormal function FIX alleles from MultiSTEP scores. Dot indicates final classifier performance metrics. **g.** Histogram depicting F9 missense variants found in the gnomAD 4.1 database according to minor allele frequencies (MAF) in hemizygotes. Color indicates functional classification based on random forest model using MultiSTEP functional scores. Vertical dashed line indicates estimated prevalence for hemophilia B in hemizygous individuals^92^. **h.** Sankey diagram of *F9* variant reinterpretation using moderate and strong levels of evidence for functional data. Labeled nodes represent the number of variants of each class.

Another important feature of hemophilia B is disease severity, defined by the level of FIX activity in plasma^20^. Severe hemophilia B is defined by undetectable FIX activity (<1%). Individuals with severe hemophilia B receive more intensive treatment and are at higher risk for complications than individuals with non-severe hemophilia B (FIX activity 1-40%)^20,21^. To explore the relationship between FIX secretion and hemophilia disease severity, we analyzed MultiSTEP secretion scores relative to disease severity reported for variants in EAHAD^23^. Variant secretion scores correlated strongly with hemophilia B severity (**Fig. 5b**). Removal of variants at epitope-adjacent positions did not appreciably affect the correlation between hemophilia B severity and MultiSTEP secretion scores (**Supplementary Fig. 10b**). Of the 258 severe disease-associated variants with secretion scores, 181 (70.2%) had low secretion scores, while 77 (29.8%) had WT-like secretion scores^23^. Many of the normally secreted variants reside at positions important for FIX activation or FIX_a_ activity including propeptide cleavage (p.R43, p.K45, and p.R46), γ-carboxylation (p.E53, p.E54, p.E72, p.E73, and p.E79), activation peptide cleavage (p.R226), or enzymatic activity (p.S411). 24 of the 77 normally secreted, severe hemophilia B-associated variants were in the propeptide or Gla domain, and 20 of these had low FIX-specific γ-carboxylation scores indicative of a defect in γ-carboxylation or Gla domain conformation (**Fig. 5c**). Three variants outside the propeptide and Gla domain also had low FIX-specific γ-carboxylation scores. The remaining 54 severe variants that scored normally in all of our assays are found throughout FIX but cluster within the heavy chain (**Fig. 5c**).

Because of the pervasive effect of cysteine substitutions on FIX secretion (**Fig. 2b,c**), we investigated their relationship to FIX antigen in plasma and disease severity. Secretion scores for gain-of-cysteine variants correlated strongly with measured FIX antigen levels in plasma (Pearson’s r = 0.88, **Supplementary Fig. 11a**). Of the nine gain-of-cysteine variants with plasma FIX antigen levels, five were undetectable, suggesting a severe phenotype. Indeed, gain-of-cysteine variants were less likely to be associated with mild disease (n = 3) compared to moderate (n = 6) or severe (n = 11) disease, although this trend was not statistically significant (two-sided Fisher’s exact test, p = 0.098; **Supplementary Fig. 11b**). Considering all substitutions, FIX antigen levels reported in EAHAD were markedly decreased for the majority of severe disease variants, compared to those causing mild or moderate disease (**Fig. 5d**). Thus, overall, loss of secretion is an important mechanism by which *F9* variants cause hemophilia B, and variants that cause very low or undetectable secretion are particularly likely to cause severe disease.

### Secretion and γ-carboxylation scores resolve FIX variants of uncertain significance

*F9* VUS are a major problem in hemophilia B, with the VUS classification almost always resulting from insufficient evidence to inform interpretation of variant pathogenicity with certainty. This evidence is lacking in part because of the rarity of the disease and in part because of very rare and novel variants discovered during sequencing^20,21^. In the MLOF genotyping program^20^, 48.6% (n = 107/220) of *F9* missense variants clinically suspected to cause hemophilia B were classified as VUS due to lack of evidence. We generated secretion and γ-carboxylation scores for 214 of these 220 MLOF variants, including 103 VUS. To resolve *F9* VUS in hemophilia B, we curated a set of 149 *F9* missense variants of known effect from ClinVar, MLOF, and gnomAD, 135 pathogenic/likely pathogenic and 14 benign/likely benign^1,3,20^. No single secretion or γ-carboxylation score set perfectly separated pathogenic and benign variants (**Fig. 5e**). Thus, we integrated the secretion and γ-carboxylation scores by developing a random forest model to classify variant function as either normal or abnormal. We trained this functional data-driven model on 111 of the 149 curated *F9* variants of known effect. The model classified 63.4% (n = 64/101) of known pathogenic and likely pathogenic training variants as abnormal function and 100% (n = 10/10) of known benign and likely benign training variants as having normal function. We then evaluated our model on the remaining 38 variants in our test set. Here, the model performed similarly well, classifying 61.8% (n = 21/34) of known pathogenic and likely pathogenic test set variants as abnormal function and 100% (n = 4/4) of known benign and likely benign test set variants as having normal function (**Fig. 5f**). When applied to all 8,528 *F9* missense variants with secretion and γ-carboxylation scores, the model classified 45.3% (n = 3,859) as functionally abnormal (**Supplementary Table 3**).

We used our functional data-driven model to annotate *F9* variants found in the gnomAD database^3^. Of 113 variants, 26 (23.0%) were annotated as abnormal function by our model (**Fig. 5g**). Of predicted abnormal function variants, p.G106S had the highest hemizygous minor allele frequency (MAF, 1.29 x 10^-5^). p.G106S is already classified as likely pathogenic and has been reported previously in multiple individuals with mild hemophilia B^20,23^. These findings are consistent with the prevalence of hemophilia B, which is estimated to affect 1 in 20,000 live male births^92^. We also evaluated our classification model’s performance across mild, moderate, and severe disease and found that it identified 47.8% of mild, 62.2% of moderate, and 75.2% of severe disease variants as abnormal function^23^ (**Supplementary Fig. 12a**). Mild and moderate disease-causing variants in the propeptide and Gla domains, which are most likely to impact γ-carboxylation-related phenotypes, are better resolved (mild: 76.5%, moderate: 72.7% severe: 80.7%, **Supplementary Fig. 12b**). Our model’s lower sensitivity to mild disease-causing variants correlates with the fact that fewer mild and moderate disease-associated variants impact FIX plasma levels (mild: 33.3%, moderate: 35.7%) as compared to severe-associated variants (56.5%) (**Fig. 5d**). 68.5% (n = 291) of gain-of-cysteine substitutions, which are poorly secreted relative to other substitutions (**Fig. 2b,c**; **Fig. 3g**), were predicted by our classification model as abnormal function, compared to 44.0% (n = 3,568) for non-cysteine variants (odds ratio: 2.75, two-sided Fisher’s exact test, p = 5.58 x 10^-23^).

Next, we compared our secretion and γ-carboxylation scores, along with the functional data-driven model, to four variant effect predictors: EVE, AlphaMissense, REVEL, and CADD^93–96^. For secretion scores, EVE (ρ = 0.611) correlated most strongly, followed by AlphaMissense (ρ = 0.545), REVEL (ρ = 0.514), and CADD (ρ = 0.413) (**Supplementary Fig. 13a**). For FIX-specific γ-carboxylation scores, AlphaMissense correlated most strongly (ρ = 0.635), followed by EVE (ρ = 0.630), REVEL (ρ = 0.601), and CADD (ρ = 0.481). A comparison of 26 deep mutational scanning datasets to 55 existing variant effect predictors found similar levels of correlations^97^.

We quantified the ability of our functional data-driven model and the four variant effect predictors to discriminate between pathogenic/likely pathogenic and benign/likely benign variants in our test set (**Supplementary Fig. 13b,c**). Both MultiSTEP and AlphaMissense had perfect positive predictive value (PPV = 1), followed by CADD (PPV = 0.94), REVEL (PPV = 0.92), and EVE (PPV = 0.71). However, MultiSTEP classified more known pathogenic/likely pathogenic variants correctly (n = 21) than AlphaMissense (n = 16). CADD and REVEL had the highest negative predictive value (NPV = 0.5), followed by MultiSTEP (NPV = 0.24), AlphaMissense (NPV = 0.18), and EVE (NPV = 0). EVE’s performance was difficult to evaluate because it did not score 31.6% (n = 12/38) of variants in our test set.

Having shown that our functional data-driven model could identify abnormal function variants, we reassessed reported *F9* missense variants classified as VUS in MLOF using evidence from the model. We applied strong evidence strength for pathogenicity to variants predicted to have abnormal function^4,20^. This resulted in reclassification of 63.1% (n = 65/103) of VUS to likely pathogenic and 67.3% (n = 66/98) of likely pathogenic variants to pathogenic (**Fig. 5h, Supplementary Table 4**). An alternative Bayesian approach suggested that our model yielded moderate evidence strength for pathogenicity^6,98,99^. Using moderate evidence strength, 61.2% (n = 63/103) of VUS were reclassified as likely pathogenic and 13.3% (n = 13/98) of likely pathogenic variants were reclassified as pathogenic (**Fig. 5h, Supplementary Table 4**). Because our classification model incorporated only secretion and γ-carboxylation functional data, it cannot identify variants that cause hemophilia B by affecting other FIX functions including direct impacts on enzymatic activity. Indeed, the model classified 38% of pathogenic variants as normal function, implying that these variants could be pathogenic because of a different type of defect (e.g. activation, catalysis, etc). Therefore, we did not use normal function predictions as evidence for benign classification. Taken together, functional evidence from MultiSTEP drastically reduced the number of *F9* VUS and lent new insight into how *F9* variants affect FIX function.

### MultiSTEP can be used to study diverse secreted proteins

Having established the utility of MultiSTEP for analyzing variant effects in FIX, we evaluated its generalizability by applying it to six additional secreted proteins: coagulation factors VII, VIII, and X, alpha-1 antitrypsin, plasma protease C1 inhibitor, and proinsulin. This diverse set of proteins ranged in size from 110 to 1,670 amino acids and included enzymes, protease inhibitors, and a peptide hormone. Antibody staining of the MultiSTEP Strep II epitope tag revealed robust surface expression in each case (**Fig. 6a**). Negative controls, including the CD28 transmembrane domain alone (i.e. without a fusion protein), deletion of the FIX start codon, and deletion of the FIX signal peptide, did not yield any appreciable signal above that found in parental cell lines. To provide further evidence that proteins displayed using MultiSTEP were properly folded, we focused on factor VIII which has been successfully expressed for therapeutic use in HEK-293 cells^100,101^. MultiSTEP-displayed factor VIII robustly bound a panel of monoclonal antibodies directed against the FVIII A1, A2, A3, C1, and C2 domains as well as two monoclonal antibodies that recognize discontinuous FVIII epitopes, confirming that displayed FVIII is folded correctly (**Fig. 6b,c**, **Supplementary Fig. 14**).

**Figure 6:**
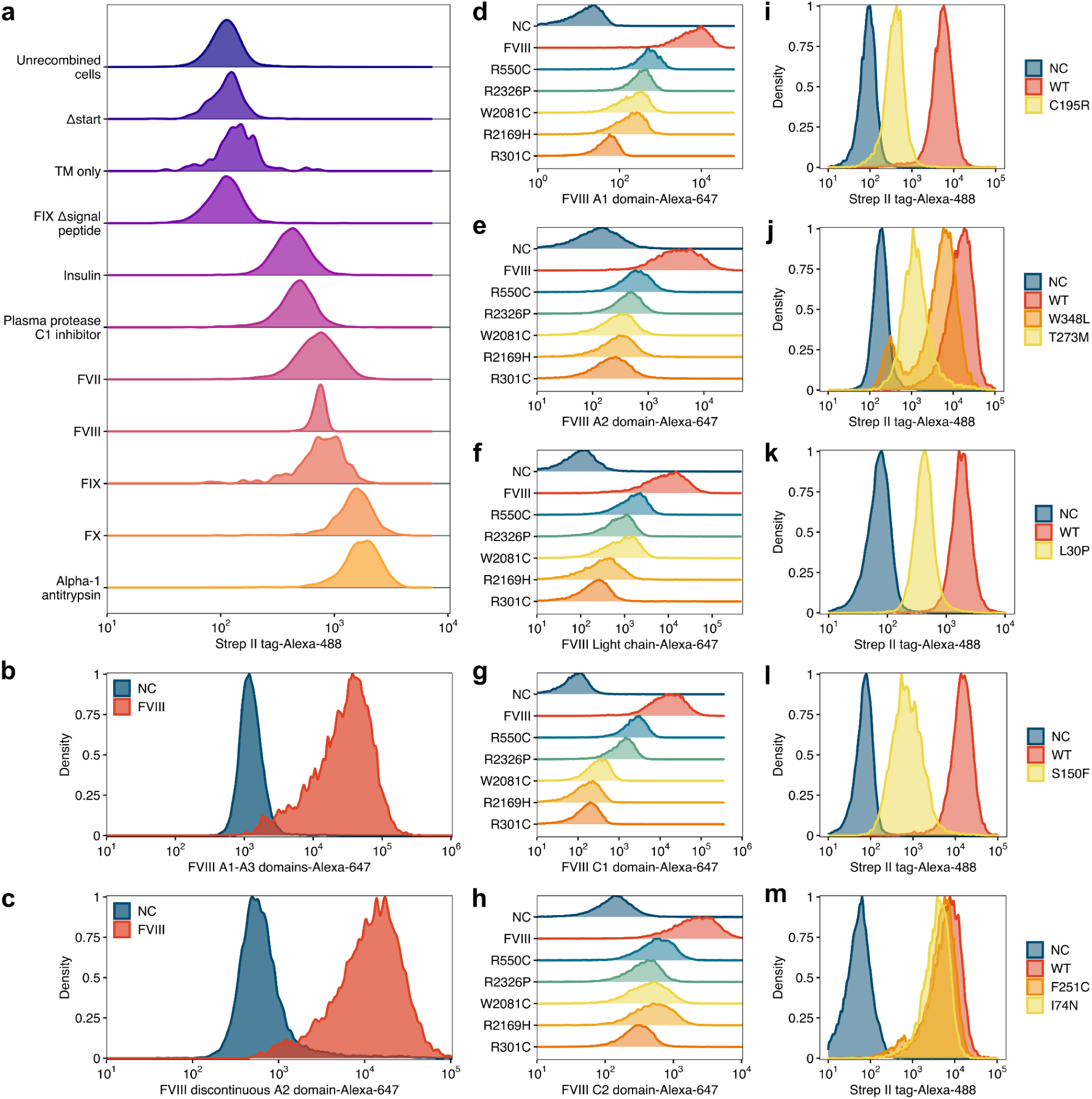
MultiSTEP can be applied to diverse secreted proteins. **a.** Flow cytometry of various protein and control constructs in the MultiSTEP backbone (n ∼30,000 cells each). Unrecombined cells do not display FIX and serve as a negative control. All other constructs contain the MultiSTEP flexible linker, strep II tag, and transmembrane domain. Δstart is a FIX cDNA that does not contain a start codon. TM only does not contain a secreted protein of interest. FIX Δsignal peptide expresses a FIX molecule without its secretion-targeting signal peptide. Fluorescent signal was generated by staining the library with a rabbit polyclonal anti-strep II tag antibody followed by staining with an Alexa Fluor-488-labeled donkey anti-rabbit secondary antibody. **b.** Flow cytometry of B-domain deleted coagulation factor VIII (FVIII) in the MultiSTEP backbone or unrecombined negative control cells (NC) (n ∼30,000 cells each). Fluorescent signal was generated by staining the library with a mouse monoclonal anti-FVIII A1-A3 antibody, which targets the discontinuous epitope at the interface of the A1 and A3 domains, followed by staining with an Alexa Fluor-647-labeled goat anti-mouse secondary antibody. **c.** Flow cytometry of B-domain deleted coagulation factor VIII (FVIII) in the MultiSTEP backbone or unrecombined negative control cells (NC) (n ∼30,000 cells each). Fluorescent signal was generated by staining the library with a mouse monoclonal anti-FVIII A2 antibody, which targets a discontinuous epitope (positions 497-510 and 584-593) within the A2 domain, followed by staining with an Alexa Fluor-647-labeled goat anti-mouse secondary antibody. . **d-h.** Flow cytometry of B-domain deleted coagulation factor VIII (FVIII) and 5 FVIII variants in the MultiSTEP backbone along with unrecombined negative control cells (NC) (n = ∼10,000 cells each). Fluorescent signal was generated by staining cells with mouse monoclonal anti-FVIII antibodies specific to the A1 (**d**), A2 (**e**), light chain (**f**), C1 (**g**), or C2 (**h**) domains, followed by staining with an Alexa Fluor-647-labeled goat anti-mouse secondary antibody. **i-m.** Flow cytometry of coagulation factor VII (**i**), coagulation factor X (**j**), proinsulin (**k**), plasma protease C1 inhibitor (**l**), and alpha-1 antitrypsin (**m**) constructs in the MultiSTEP backbone along with unrecombined negative control (NC) (n ∼10,000 cells each). For each secreted protein, at least one variant with clinical or *in vitro* evidence of decreased secretion is included. Fluorescent signal was generated by staining the library with a rabbit polyclonal anti-strep II tag antibody followed by staining with an Alexa Fluor-488-labeled donkey anti-rabbit secondary antibody.

We tested 5 FVIII variants known to impact FVIII secretion and found that we could identify a wide range of secretion defects using several different antibodies (**Fig. 6d-h**). For the other five secreted proteins, we identified at least one variant from public databases (EAHAD, LOVD) or individual publications that had either clinical (plasma antigen) or *in vitro* evidence of decreased secretion^102–106^. Variants in FVII, FX, insulin, and plasma protease C1 inhibitor all demonstrated a marked decrease in surface expression (**Fig. 6i-l**), as expected. For variants in alpha-1 antitrypsin, we observed a small decrease in secretion (two sided t-test, *p* = 0.046 and *p* = 0.049, **Fig. 6m**).

## Discussion

MultiSTEP is a generalizable, multiplexed method for measuring the effects of variants on the function of human secreted proteins. MultiSTEP can be combined with diverse antibodies to quantify secretion and post-translational modifications, which until now have been beyond the reach of MAVEs. We applied MultiSTEP to coagulation factor IX, a protein critical for hemostasis, measuring 44,816 variant effects on secretion and post-translational γ-carboxylation for 8,528 missense and 436 synonymous variants using a panel of five antibodies. We revealed that nearly half of FIX missense variants caused some degree of loss of function, mostly by reducing or eliminating secretion. In particular, unlike for intracellular proteins, either adding or removing cysteines profoundly impacted FIX secretion, likely reflecting the importance of correct disulfide bonding on FIX structure^33,40,41,58^.

Most variants located outside of the FIX propeptide and Gla domain did not show effects with either carboxylation-sensitive antibody, supporting the notion that there is not another FIX domain important in FIX γ-carboxylation. Propeptide and Gla domain variants with reduced scores for one or both carboxylation-sensitive antibodies also generally did not impact FIX secretion. In the propeptide, our results suggest that variants that affect propeptide cleavage can alter Gla-domain conformation while permitting the γ-carboxylation of at least some glutamates in the conserved Gla-domain motif^28,81,82^. In the Gla domain, glutamates at positions known to be γ-carboxylated do not all appear to play the same role, with those that coordinate calcium or comprise the C-terminal aromatic stack being markedly more sensitive to variation than those that generally coordinate magnesium, consistent with the importance of calcium coordination for core Gla domain structure and function^23,83,84,86,107^.

Genetic variation in *F9* is the primary cause of hemophilia B, and pathogenic missense FIX variants can disrupt protein secretion or function. Variant secretion correlated strongly with circulating FIX antigen levels in people. More than 40% of variants associated with severe disease had profound secretion defects, illustrating the importance of loss of secretion as a causal mechanism of hemophilia B. We developed a random forest classifier to unify the multiple secretion and γ-carboxylation scores we collected for each variant in order to provide evidence for interpreting significance of *F9* variants. Using the results of this functional data-driven model, we reevaluated the classifications of all *F9* missense variants found in the MLOF hemophilia genotyping program. Evidence from the model enabled us to upgrade 65 of 103 *F9* missense VUS to likely pathogenic and 66 likely pathogenic variants to pathogenic. These model predictions are available for 8,528 (97.4%) FIX missense variants, meaning that newly discovered variants in hemophilia B can be more accurately and quickly classified in the future. The American College of Medical Genetics and Genomics (ACMG) guidelines indicate that variant effect predictors and functional data are independent lines of evidence and should be combined when classifying a variant^4^. We also showed that several variant effect predictors could distinguish pathogenic and benign variants. Thus, the high-quality, comprehensive *in vitro* functional data we furnish, along with variant effect predictions, promise to dramatically diminish the number of VUS in *F9* going forward.

MAVE-compatible *E. coli* and *S. cerevisiae* cell surface display methods have typically been applied to intracellular proteins^108–111^. Moreover, these methods are not generally suitable for human secreted proteins which require specialized intracellular trafficking pathways and extensive post-translational modification. Mammalian cell surface display systems that allow for the study of post-translational modifications rely on non-native signal peptides^112–114^. The use of alternative or artificial signal peptides prevents accurate assessment of the functional effects of native signal peptide variants and can also disrupt protein folding, trafficking, localization, or post-translational modification^42,43,115^. Approaches to display IgG antibodies using endogenous signal peptides have been developed recently, but, unlike MultiSTEP, rely on transient transfection or lentiviral integration to express variants^112,116,117^. MultiSTEP avoids common issues with these approaches that can obscure variant functional effects, including lentiviral template switching, variability in transgene expression from random integration sites, and expression of multiple variants per cell^118,119^. Instead, MultiSTEP employs a genomically integrated, multiplexable recombinase-based landing pad system we developed and have used extensively to study non-secreted proteins^31,32^. This landing pad system ensures integration of a single variant per cell, avoiding many of the issues with other methods. However, integrating a landing pad into a cell line and validating it is more laborious than other methods. Thus, by enabling multiplexed human cell surface display of secreted protein variants, MultiSTEP meets a unique and important need.

While our FIX variant secretion and γ-carboxylation results illuminate various structural and biochemical features of FIX, identify pathogenic variant mechanisms, and improve clinical variant interpretation, MultiSTEP has limitations. For example, while all seven of the secreted proteins we displayed using MultiSTEP were expressed on the cell surface, the folding, secretion or function of some other proteins may be compromised by linkage to the C-terminal transmembrane domain fusion. The expression of protein on the surface could be affected by the choice of transmembrane domain, and some proteins might not be amenable to display or may require display in a particular cell type. We chose the 293-F cell line because it has been used to produce a wide variety of human proteins, improves protein expression, and enables large-scale culture. The 293-F cell line also grows in suspension, eliminating potential difficulties with dissociating adherent cells^15,100,113,117,120–125^. MultiSTEP could be implemented in other cell lines where recombinase-based landing pads work, including HeLa, NIH3T3, HEK293T, HAP-1, A549, HEK293FT, K562, HT29, HCT116, U2OS, HDFa, COS-1, Jurkat, and CHO cell lines as well as in hESCs, iPSCs, and mouse embryonic fibroblasts^31–33,41,58,126–142^. While we demonstrated a MultiSTEP-compatible non-enzymatic cell dissociation method, this method might adversely impact the integrity of displayed proteins.

Using antibodies as detection reagents also imposes limitations. For example, variants that disrupt antibody epitopes can give falsely low signals so we recommend assaying variants with multiple antibodies. Some antibodies, like the FIX-specific carboxylation-sensitive antibody we used, require folding for binding, and these requirements must be accounted for when interpreting the results. Antibody quality is also important. The Strep II tag antibody assay replicates were less well correlated than the other antibody assays. Since all assays were executed with the same library and with similar parameters (e.g. number of cells sorted, depth of sequencing, analysis pipeline, etc), we believe the Strep II tag antibody assay itself was the cause of the difference in data quality. Thus, selecting performant antibodies is critical. We suggest evaluating and validating candidate antibodies carefully, and, where feasible, comparing the results of multiple antibodies^143^. Lastly, no functional assay fully recapitulates *in vivo* biology. For example, we did not measure FIX enzymatic activity in our system. Therefore, we discourage using our results as evidence for benign variant classification in hemophilia B.

MultiSTEP is a generalizable platform for measuring the effects of missense variation in secreted proteins, complementing the plethora of MAVEs already developed to study intracellular proteins. Because MultiSTEP is compatible with reagents that produce a fluorescent signal, we were able to probe multiple functional aspects of FIX using a panel of antibodies. Antibodies are available for numerous clinically and biologically important secreted proteins, as well as a variety of post-translational modifications. Moreover, other fluorescently labeled reagents such as protein binding partners or labeled covalent substrates could be used to read out other functions. Lastly, cells could be interrogated for the presence of intracellular proteins, or fragments thereof, offering the opportunity to measure the abundance of variants both inside and outside the cell. Such measurements could offer insights into mechanisms leading to poor secretion such as altered folding, stability, or trafficking. The ability to characterize the diverse effects of massive numbers of secreted human protein variants on the surface of human cells creates the opportunity to understand variant effects on secreted protein structure, function, and pathogenicity.

## Methods

### General reagents

Chemicals were purchased from ThermoFisher Scientific and synthetic oligonucleotides were purchased from IDT unless otherwise noted. Sequences of oligonucleotides used in this work can be found in **Supplementary Table 5**.

All *E. coli* were grown in Luria Broth (LB) at 37°C with shaking at 225 rpm for 16-18 hours with 100 μg/mL carbenicillin, unless otherwise indicated. Routine cloning was performed in homemade chemically competent Top10F’ *E. coli*, whereas library cloning was performed in commercially available electrocompetent NEB-10β *E. coli* (New England Biolabs).

Inverse PCR reactions, unless otherwise specified, were performed in 30 μL reactions with 2x Kapa HiFi ReadyMix (Kapa Biosystems) or Q5 2x master mix (New England Biolabs), with 40 ng starting plasmid and 0.15 μM each forward and reverse primers. Reaction conditions were 95°C for 3 minutes, 98°C for 20 seconds, 61°C for 30 seconds, 72°C for 1 minute/kb, repeating for 8 total cycles, then followed by a final 72°C extension for 1 minute/kb, and held at 4°C. PCR products were then digested at 37°C with DpnI (New England Biolabs) for 2 hours to remove residual starting plasmid, followed by heat inactivation at 80°C for 20 minutes. PCR products were then checked on a 1% agarose gel with 1x SYBR Safe at 100 V, 45 minutes for purity and size, and gel extracted if needed.

For large modifications (>50 bp), PCR products were then Gibson assembled using a 3:1 molar ratio of insert(s):backbone at 50°C for 1 hour, after which 2 μL of product was transformed into homemade chemically competent Top10F’ *E. coli*^144^. For small modifications, *in vivo assembly* cloning (IVA cloning) of linear products was used by transforming 5 μL of PCR product directly into Top10F’ *E. coli* without recircularization^145^.

For both Gibson assembly and IVA cloning, after addition of PCR-amplified DNA, Top10F’ cells were incubated on ice for 30 minutes before a 30 second heat shock at 42°C. Cells were then returned to ice for 2 minutes before being added to 1 mL SOC to recover for 1 hour at 37°C, shaking at 225 rpm. After recovery, 100 μL was spread on LB-ampicillin plates and grown overnight for 16-18 hours. Colonies were then screened for correct insertion by Sanger sequencing (small modifications) or colony PCR (large modifications), Sanger sequence confirmed, and isolated using miniprep or midiprep kits according to the manufacturer’s instructions (Qiagen).

A Golden Gate assembly-compatible MultiSTEP vector (MultiSTEP-GG) was designed with BsmBI cut sites flanking the ORF. For Golden Gate assembly of each construct, 75ng of the MultiSTEP-GG plasmid were combined with a 2:1 molar ratio of each gene fragment. DNA was incubated in a 25μL Golden Gate reaction containing 10x T4 ligase buffer, 3μL Esp3I, and 2.5μL T4 DNA ligase (both enzymes from NEB). The Golden Gate reaction was incubated on a thermocycler for 5 minutes at 37°C followed by 5 minutes at 16°C. After 30 cycles, there was a final cycle of 15 minutes at each temperature, followed by heat inactivation at 85°C. To decrease background, an additional 3μL water, 1μL Esp3I, and 1μL 10x CutSmart buffer were added to each reaction mixture and incubated for four hours at 37°C. The reaction mix was cleaned using a NEB Clean and Concentrate kit, then transfected into chemically competent bacteria using standard protocols.

### Variant nomenclature

Variants in FIX and other proteins are described using the Human Genome Variation Society (HGVS) numbering system, which designates the start position (1) of proteins as the first methionine of their dominant isoform^146^. For FIX, multiple other variant numbering systems have been used widely. In particular, the legacy system (based on mature FIX protein) and the chymotrypsin system (based on evolutionary similarity of serine proteases like FIX to chymotrypsin) are widely used, especially in older publications. Direct conversions between the HGVS, legacy, and chymotrypsin numbering systems for FIX can be found in **Supplementary Table 6**.

### Cloning into the MultiSTEP landing pad donor plasmid

To clone attB-F9-10L-strepII-10L-CD28-IRES-mCherry (pNP0001), first, an inverse PCR was performed on attB-EGFP-PTEN-IRES-mCherry-562bgl with primers NP0207 and NP0325 to remove EGFP-PTEN and create compatible Gibson overhangs using Kapa HiFi polymerase (Kapa Biosystems). A gBlock (NPg0007) containing human *F9* cDNA and a second gBlock (NPg0012) containing a GC-optimized 10 amino acid (GGGGS)_2_ flexible linker, a strep II protein tag, and the single-pass transmembrane domain of CD28 was then assembled following the standard Gibson assembly cloning protocol above. After sequence-confirmation, a second round of inverse PCR was performed to insert a second (GGGGS)_2_ flexible linker after the strep II protein tag using primers NP0334 and NP0356 following the standard IVA cloning protocol above.

To clone an empty tethering backbone vector attB-10L-strepII-10L-CD28-IRES-mCherry (pNP0079), *F9* was removed by IVA cloning following the same protocol as above with primers NP0325 and NP0377. This plasmid was used as Δstart in **Fig. 6**. To remove the *F9* signal peptide (pNP0010), primers NP0295 and NP0471 were used following the standard IVA cloning protocol above. To create a transmembrane-only plasmid (pNP0009), primers NP0325 and NP0470 were used following the standard IVA protocol above.

To clone missense *F9* variants for assay validation, single amino acid variants were introduced into pNP0001 by IVA cloning following the same protocol as above. For each variant, the following primers were used: p.C28Y (pNP0011): NP0488 and NP0489; p.A37T (pNP0003): NP0293 and NP0294; p.G58E (pNP0027): NP0225 and NP0226; p.E67K (pNP0019): NP0217 and NP0218; p.C134R (pNP0080): NP0591 and NP0592; p.S220T (pNP0057): NP0478 and NP0479; p.H267L (pNP0065): NP0273 and NP0274.

cDNA constructs for human *F7*, *F10*, *SERPINA1*, *SERPING1*, and *INS* were ordered from the Mammalian Gene Collection (Horizon Discovery) and cloned into the landing pad donor backbone (pNP0079) using Gibson assembly. For each gene, the donor backbone was amplified using NP0295 and NP0325. To generate pNP0088, *F7* cDNA was amplified with NP0615 and NP0616. To generate pNP0089, *F10* cDNA was amplified with NP0617 and NP0618. To generate pNP0090, *SERPINA1* cDNA was amplified with NP0619 and NP0620. To generate pNP0091, *SERPING1* cDNA was amplified with NP0621 and NP0622. To generate pNP0092, *INS* cDNA was amplified with NP0623 and NP0624.

pcDNA4/Full Length FVIII was a gift from Robert Peters (Addgene #41036). To generate attB-F8-10L-StrepII-10L-CD28-IRES-mCherry (pNP0084), an inverse PCR was performed on pcDNA4/Full Length FVIII using NP0398 and NP0399 to isolate *F8.* pNP0079 backbone was also inverse PCR amplified using NP0400 and NP0401 to add *F8* overhangs. The products were Gibson assembled using the standard cloning protocol above.

Because full length *F8* is poorly expressed both in cell culture and *in vivo*, we adapted a B-domain deletion version of *F8* cDNA shared by Dr. Steven Pipe (University of Michigan) that has improved coagulation factor VIII expression and retains FVIII function^147–149^. Primers NP0405 and NP0406 were used to create B-domain deleted *F8* (pNP0085) using the IVA cloning protocol above.

To clone variants for *F7* (p.C195R), *F8* (p.R301C, p.R550C, p.W2081C, p.R2169H, and p.R2326P), *INS* (p.L30P), and *SERPING1* (p.S150F), variant allele gene fragments were ordered from Twist Bioscience. Gene fragments were resuspended in TE elution buffer and cloned into the MultiSTEP-GG vector following the cloning protocol above. To clone variants in *F10* and *SERPINA1*, the following primers were used with the IVA cloning protocol above: *F10* p.T273M: KL01 and KL02; *F10* p.W348L: KL03 and KL04; *SERPINA1* p.I74N: KL05 and KL06; *SERPINA1* p.F251C: KL07 and KL08.

### Site-saturation mutagenesis library cloning

Site-saturation mutagenesis oligonucleotide pools were ordered from Twist Biosciences for each position in *F9* (**Supplementary Table 1**). Each position contained one codon for each synonymous or missense variant. Thus, our library was designed to include all 8,759 possible missense variants and 461 synonymous variants, one for each position (**Supplementary Table 1**). 50 ng of each oligonucleotide were resuspended in 10 μL water, and then pooled in equal volumes into three tiled sublibraries encompassing the entire length of *F9*, including 20 positions of overlap between adjacent sublibraries. Tile 1 sublibrary: positions p.Q2-p.K164; Tile 2 sublibrary: positions p.A146-p.I318; Tile 3 sublibrary: positions p.L299-p.T461.

pNP0079 was inverse PCR amplified using NP0295 and NP0325 following the standard protocol, and the correct size PCR product was gel extracted. The backbone PCR product was then Gibson assembled with each of the three pooled sublibraries at a 5:1 molar ratio of insert:backbone at 50°C for 1 hour. Gibson assembled products were then cleaned and eluted in 10 μL water (Zymo Clean and Concentrate). 1 μL of cleaned product was then added to 25 μL NEB-10β *E. coli* in pre-chilled cuvettes and allowed to rest on ice for 30 minutes. 2 electroporation replicates were performed per sublibrary tile, as well as a pUC19 control (10 pg/μL). Cells were then electroporated at 2 kV for 6 milliseconds. After electroporation, cells were immediately resuspended in 100 μL pre-warmed SOC and transferred to a culture tube. Identical replicates were pooled at this step, and pre-warmed SOC was added to a final volume of 1 mL and allowed to recover at 37°C, shaking at 225 rpm, for 1 hour. After recovery, the entire recovery volume was added to 49 mL of LB containing 100 μg/mL carbenicillin and allowed to grow overnight. After 2-3 minutes of shaking, a 200 μL sample was taken and used for serial dilutions to estimate colony counts on LB-ampicillin plates as a proxy for the number of unique molecules transformed and to gauge coverage of the library. After 16 hours of overnight growth at 37°C, each 50 mL midiprep culture was spun down for 30 minutes at 4,300 x *g* and plasmid DNA isolated using a midiprep kit according to the manufacturers instructions (Qiagen).

### Barcoding site-saturation mutagenesis libraries

To barcode each sublibrary, 1 μg of each sublibrary plasmid was digested at 37°C for 5 hours with NheI-HF and SacI-HF (New England Biolabs), incubated with rSAP for 30 minutes at 37°C, then heat-inactivated at 65°C for 20 minutes. Digested product was then run on a 1% agarose with 1x SYBR Safe (ThermoFisher Scientific) gel for 45 minutes at 100 V and gel extracted (Qiagen).

An IDT Ultramer (NP0490) with 18 degenerate nucleotides was resuspended at 10 μM. 1 μL NP0490 was then annealed with 1 μL of 10 μM NP0397 primer, 4 μL CutSmart buffer, and 34 μL water by running at 98°C for 3 minutes, followed by ramping down to 25°C at −0.1°C per second. After annealing, 1.35 μL of 1 mM dNTPs and 0.8 μL Klenow exo-polymerase (New England Biolabs) were added to fill in the barcode oligo. The cycling conditions were 25°C for 15 minutes, 70°C for 20 minutes, and then ramped down to 37°C in −0.1°C per second increments. Once at 37°C, 1 μL each NheI-HF and SacI-HF were added and digested for 1 hour. Digested product was then run on a 4% agarose gel with 1x SYBR Safe for 45 minutes at 100 V and gel extracted (Qiagen).

Both gel extracted sublibrary and barcode oligonucleotide were cleaned and eluted in 10 μL and 30 μL water, respectively (Zymo Clean and Concentrate). A 7:1 molar ratio of barcode oligonucleotide to sublibrary was ligated overnight at 16°C with T4 DNA ligase (New England Biolabs).

Ligated product was then cleaned and eluted in 10 μL water (Zymo). 1 μL of ligation product, ligation controls, or pUC19 was electroporated into NEB-10β *E. coli* following the same procedure as above. For sublibrary ligation products, 2 independent replicates were pooled before recovery. After recovery, each sublibrary ligation product was bottlenecked by diluting various recovery volumes (500 μL, 250 μL, 125 μL, and 50 μL) into independent 50 mL LB-ampicillin cultures. After 2-3 minutes of shaking, a 200 μL sample from each ligation bottleneck was taken and used for serial dilutions to estimate colony counts on LB-ampicillin plates. Colony counts were then used to estimate the number of barcoded variants present in each sublibrary for each bottleneck. After 16 hours of overnight growth at 37°C, each 50 mL midiprep culture was spun down for 30 minutes at 4,300*g* and plasmid DNA isolated using a midiprep kit according to the manufacturer’s instructions (Qiagen).

### Estimation of barcoded variant coverage by Illumina sequencing

Each barcoded and bottlenecked plasmid sublibrary was diluted to 10 ng/μL and amplified for Illumina sequencing to more accurately determine the number of unique barcodes present. Briefly, to add adapter sequences, primers NP0492 and NP0493 were mixed at a final concentration of 0.5 μM with 10 ng plasmid DNA, 25 μL Q5 polymerase (New England Biolabs), and 19 μL water. Cycling conditions were an initial denaturing at 98°C for 30 seconds, followed by 5 cycles of 98°C for 10 seconds, 61°C for 30 seconds, and 72°C for 30 seconds, followed by a final 72°C extension for 2 minutes and a hold at 4°C. PCR products were then cleaned using 0.8x AmpureXP beads (Beckman Coulter) and eluted in 16 μL water following the manufacturer’s instructions.

The entire elution volume for each sample was then mixed with 25 μL Q5 polymerase, 0.25 μL of 100x SYBR Green I, 4.75 μL water, and one uniquely indexed forward (NP0565-NP0578) and reverse (NP0551-NP0564) primer at a final concentration of 0.5 μM. Samples were run on the CFX Connect (Bio-Rad) for a maximum of 15 cycles or until all samples were above 3,000 relative fluorescence units. Reactions were denatured at 98°C for 30 seconds, and cycled at 98°C for 10 seconds, 65°C for 30 seconds, and 72°C for 30 seconds, with a final extension at 72°C for 2 minutes. Samples were then run on a 2% agarose gel with 1x SYBR Safe for 2 hours at 120 V before gel extraction with a Freeze ‘N Squeeze column (Bio-Rad). After extraction, samples were quantified using the Qubit dsDNA HS Assay Kit (ThermoFisher Scientific), pooled in equimolar concentrations, and sequenced on a NextSeq 550 using a NextSeq 500/550 High Output v2.5 75 cycle kit (Illumina) using custom sequencing primers NP0494-NP0497. Using a custom script, sequencing reads were converted to FASTQ format and demultiplexed using bcl2fastq (v2.20), forward and reverse barcode reads were paired using PEAR (v0.9.11), and unique barcodes were counted and filtered^150^.

### PacBio sequencing of FIX sublibraries for variant-barcode mapping

2.5 μg of each library subtile was digested with AflII (New England Biolabs) in CutSmart buffer for 4 hours at 37°C, followed by heat inactivation at 65°C for 20 minutes and purified with AMPure PB beads (Pacific Biosciences, 100-265-900). All library preparation and PacBio DNA sequencing were performed at University of Washington PacBio Sequencing Services. At all steps, DNA quantity was checked with fluorometry on the DS-11 FX instrument (DeNovix) with the Qubit dsDNA HS Assay Kit (ThermoFisher Scientific) and sizes were examined on a 2100 Bioanalyzer (Agilent Technologies) using the High Sensitivity DNA Kit. SMRTbell sequencing libraries were prepared according to the protocol “Procedure & Checklist - Preparing SMRTbell® Libraries using PacBio® Barcoded Universal Primers for Multiplexing Amplicons” and the SMRTbell Express Template Prep Kit 2.0 (Pacific Biosciences, 100-938-900) with barcoded adapters (Pacific Biosciences, 101-628-400).

After library preparation, the barcoded libraries were pooled by normalizing mass to the number of constructs contained in each pool. The final library was bound with Sequencing Primer v4 and Sequel II Polymerase v2.0 and sequenced on two SMRT Cells 8M using Sequencing Plate v2.0, diffusion loading, 1.5 hour pre-extension, and 30-hour movie time. Additional data were collected after treatment with SMRTbell Cleanup Kit v2 to remove imperfect and damaged templates, using Sequel Polymerase v2.2, adaptive loading with a target of 0.85, and a 1.3 hour pre-extension time. CCS consensus and demultiplexing were calculated using SMRT Link version 10.2 with default settings and reads that passed an estimated quality filter of ≥Q20 were selected as “HiFi” reads and used to map barcodes to variants.

“HiFi” PacBio reads were first subjected to a custom analysis pipeline, AssemblyByPacBio. Each consensus CCS sequence was aligned to the WT *F9* cDNA sequence using BWA-MEM v0.7.10-r789, generating CIGAR and MD strings, which are then used to extract barcodes and the variable region containing *F9*^151^. The output from AssemblyByPacBio was then passed through PacRAT, which takes all CCS reads containing the same barcode and performs multiple sequence alignment to improve variant calling^152^. Alignments with a variant agreement threshold of less than 0.6 or fewer than 3 independent CCS reads were filtered from further analysis, resulting in 260,224 unique barcodes across all three sublibraries. A custom R script was then used to parse the PacRAT output and generate a final barcode-variant map, containing 8,528 missense and 439 synonymous variants (**Supplementary Table 1**).

### General cell culture conditions

HEK293T cells (ATCC CRL-3216) were grown at 37°C and 5% CO_2_ in Dulbecco’s modified Eagle’s medium (ThermoFisher Scientific) supplemented with 10% fetal bovine serum (ThermoFisher Scientific), 100 U/mL penicillin, and 100 ng/mL streptomycin (ThermoFisher Scientific). Cells were passaged every 2-3 days by detachment with 0.05% trypsin-EDTA (ThermoFisher Scientific).

Freestyle 293-F cells were grown in Freestyle 293 Expression Medium (ThermoFisher Scientific) at 37°C and 8% CO_2_ while shaking at 135 rpm. Cells were regularly counted using trypan blue staining on a Countess II FL automatic hemocytometer (ThermoFisher Scientific) and passaged by dilution to 3 x 10^5^ cells/mL once reaching a concentration between 1 x 10^6^ and 2 x 10^6^ cells/mL, unless otherwise stated. All Freestyle 293-F cells containing a landing pad were induced with 2 μg/mL doxycycline (Sigma-Aldrich).

### Lentiviral transduction to generate suspension Freestyle 293-F landing pad line

To generate a landing pad lentiviral vector, 2.5 x 10^5^ cells HEK293T cells were passaged into 6 well plates, and transfected with 500 ng pMD-VSVg (Addgene #12259), 1,750 ng psPax2 (Addgene #12260), and 1,750 ng landing pad G384A vector template using 6 μL Fugene 6 (Promega) following the manufacturer’s protocol. The next day, media was exchanged, and the supernatant was collected every 12 hours for the following 72 hours to harvest lentivirus. The supernatant was then centrifuged at 300 x *g* for 10 minutes and passed through a 0.45 μm filter, before being aliquoted, flash frozen in liquid nitrogen, and stored at −80°C^31^.

1 x 10^7^ Freestyle 293-F cells were plated in 20 mL media and then incubated with varying volumes of lentivirus-containing supernatant (1 mL to 1 μL). 24 hours later, media was removed and cells were washed once before replating into 30 mL media. On day 4 post-transduction, 2 μg/mL doxycycline was added to the cells, which were then grown for 10 more days with regular passaging.

Cells were then washed with PBS + 1% bovine serum albumin (BSA, Sigma-Aldrich) before assessing mTagBFP2 fluorescence from the landing pad on an LSR II (BD Biosciences). Only samples with a multiplicity of infection (MOI) <1 were then sorted by FACS on an Aria III for mTagBFP2+ cells. A total of 17,114 BFP+ cells were replated into a half deep 96 well plate (Applikon Biotechnology) and allowed to expand with the addition of 2 μg/mL doxycycline and 100 μg/mL blasticidin (Invivogen) to select for functional landing pad cells. Single landing pad integration was confirmed by co-transfection of EGFP and mCherry recombination vectors.

### Fluorescence-activated cell sorting parameters

For all experiments, the following settings and gates were used to sort individual cells. Live, cells were first identified using FSC-A vs SSC-A. Live cells were then gated for single cells using two sequential gates–the first: FSC-A vs FSC-H, and the second: SSC-A vs SSC-H. mTagBFP2 expression from the unrecombined landing pad was excited using the 405 nm laser and captured on 450/50 nm bandpass filter. A 410 nm long pass filter preceded the 450/50 nm bandpass filter on the Symphony A3. mCherry expressed from the recombined landing pad was excited using the 561 nm laser and captured using a 595 nm (LSR II) or 600 nm (Aria III and Symphony A3) long pass and 610/20 nm bandpass filters. EGFP or Alexa488-labeled antibodies were excited using the 488 nm laser and captured using a 505 nm long pass and 530/30 nm bandpass filters (LSR II and Aria III) or a 515/20 bandpass filter (Symphony A3). Alexa647-labeled antibodies were excited using a 637 nm (LSR II and Symphony A3) or 640 nm laser (Aria III) and captured using a 670/30 nm bandpass filter (LSR II and Aria III) with an additional 650 nm long pass filter for the Symphony A3. All flow cytometry data were collected with FACSDiva v.8.0.1 and analyzed using FlowJo v.10.7.1.

### Recombination of Freestyle 293-F cells

Freestyle 293-F cells were transfected at 1 x 10^6^ cells/mL with 293Fectin following the manufacturer’s protocol with the following alterations. Briefly, for every 1 mL of cells transfected, 2 μL 293Fectin was mixed with 31.5 μL OPTI-MEM in one tube, and 1 μg of total plasmid DNA was added to OPTI-MEM for a final volume of 33.5 μL in a second tube. For recombination experiments, the 1 μg of total DNA was split in a 1:15 ratio of pCAG-NLS-Bxb1 (Addgene #51271) and recombination vector. After 5 minutes at room temperature, the two tubes were mixed together by gentle pipetting and incubated at room temperature for 20 minutes, before being added to cells. For single variants or controls, 1 x 10^7^ cells in 10 mL were transfected. For libraries, 3 x 10^7^ cells in 30 mL of cells were transfected.

48 hours after transfection, cells were split 1:9 into two separate flasks with 2 μg/mL doxycycline. The second flask containing 9 parts was additionally treated with 10 nM rimiducid (AP1903) to selectively kill off unrecombined cells. Two days after rimiducid treatment, cells were counted using trypan blue exclusion, and then live cells were separated from dead cells using Histopaque-1077 (Sigma-Aldrich). Cells were diluted to 35 mL total volume and then layered slowly on top of 15 mL Histopaque-1077 in a 50 mL conical. The cells were then centrifuged at 400 x *g* for 30 minutes with no acceleration and no break. The top 30 mL of media was aspirated, and the cells at the interface between Histopaque-1077 and media were removed and resuspended in 30 mL final volume of media. These cells were centrifuged again at 300 x *g* for 5 minutes, the media was removed, and the pellet was resuspended in 30 mL of fresh media and 2 μg/mL doxycycline. Cells were counted using trypan blue to determine yield. Approximately 1 week after transfection, cells were assayed on an LSR II or Symphony A3 for mCherry fluorescence to determine recombination rate and selection efficiency. Cells were maintained with 2 μg/mL doxycycline throughout.

### ELISA

Constructs containing FIX variants and mCherry reporter were recombined into Freestyle 293-F cells and isolated as above. Cells were allowed to grow in the absence of doxycycline before seeding to approximately equal numbers of mCherry-positive cells per flask. Expression of FIX was induced via doxycycline, and supernatant samples were taken at 48 hours.

FIX abundance in WT-expressing cell supernatants was calculated for each time point relative to pooled normal plasma (Precision Biologic) using a FIX matched-pair polyclonal antibody EIA kit (Enzyme Research Labs). The 48 hour time point supernatants were assayed for FIX abundance relative to timepoint-matched WT FIX supernatant over multiple assays containing four dilutions in duplicate for each supernatant. The FIX concentration for each well was interpolated from the standard curve for each plate. Values outside the linear range of the curve were discarded. Mean FIX abundance in supernatant was calculated from the pool of in-range values across all assays and replicates for each sample and time point.

### Antibody staining for surface-displayed proteins

For all secretion antibodies, cold PBS + 1% BSA was used as a staining and dilution buffer. Because proper folding of the γ-carboxylated Gla domain is dependent on calcium, a 1:10 dilution of cold PBS + Ca/Mg + 1% BSA into PBS + 1% BSA was used as a staining and dilution buffer for γ-carboxylation antibodies. See **Supplementary Table 7** for additional antibody details, stock concentrations, and working dilutions.

Cells were plated at 3 x 10^5^ cells/mL in either 10 mL (single variants and controls) or 30 mL (libraries) of Freestyle media with the addition of 50 nM vitamin K_1_ (Sigma-Aldrich). For experiments involving γ-carboxylation antibodies, one control flask of WT FIX cells was additionally supplemented with 100 nM warfarin (Sigma-Aldrich) to reduce carboxylation of the FIX Gla domain. After 24 hours, cells were induced with 2 μg/mL doxycycline and grown for an additional 48 hours.

On the day of staining, flasks of cells were split into equal 4 mL volumes (6 per 30 mL flask, 1 per 10 mL single variant or control) and spun at 300 x *g* for 3 minutes. Media was aspirated, and 3 washes of 1 mL cold staining buffer were performed, with 300 x *g* spins and supernatant aspiration between. Cells were then resuspended in 100 μL of diluted primary antibodies and incubated at room temperature for 30 minutes, with vortexing at 10 minute intervals. After primary antibody staining, cells were diluted with 1 mL staining buffer and spun at 300 x *g* for 3 minutes. This washing step was repeated twice more before secondary antibody staining. For secondary antibody staining, cells were resuspended in 100 μL of diluted secondary antibodies and incubated at room temperature for 30 minutes in the dark, with vortexing at 10 minute intervals. After secondary antibody staining, cells were again diluted with 1 mL cold staining buffer and spun at 300 x *g* for 3 minutes. This washing step was repeated twice more before a final resuspension in 500 μL staining buffer. At this point, all identical tubes were pooled.

For experiments not requiring FACS, cells were analyzed by flow cytometry for antibody fluorescence on either the LSR II or Symphony A3 cytometers. For library sorts, FACS was performed on an Aria III, dividing the library into four approximately equally sized quartile bins based on the ratio of fluorescent antibody to mCherry fluorescence. At least 2 million cells were sorted into each bin.

After sorting, each bin of library cells was spun down at 300 x *g* for 5 minutes and then resuspended in 10 mL of Freestyle 293 Expression media and supplemented with 100 U/mL penicillin and 100 ng/mL streptomycin. Cells were counted daily for 4-6 days, diluting to 20 mL final volume once the concentration was above 5 x 10^5^ cells/mL. Cells were harvested once each bin contained at least 20 million cells. To harvest, cells were spun down at 300 x *g* for 10 minutes, the supernatant was aspirated, and the pellet was flash frozen in liquid nitrogen before storage at −20°C.

### Genomic DNA prep, barcode amplification, and sequencing

Genomic DNA was prepared according to previously described protocols^33,41,58^. Briefly, genomic DNA was extracted from harvested cell pellets using the DNEasy Blood and Tissue kit (Qiagen) according to the manufacturer’s protocol with the addition of a 30 minute RNase digestion at 56°C during the resuspension step. Six DNEasy columns were used per cell pellet.

Two technical PCR replicates were performed on each bin for each sample to assess the variability in barcode counts from sequencing and amplification (mean r = 0.95, **Supplementary Table 8**). The protocol outlined above (Estimation of barcoded variant coverage by Illumina sequencing section) was used to amplify each sample with the following changes: 1) For each replicate, 8 identical 50 μL first-round PCRs were prepared, and each PCR contained 2,500 ng genomic DNA, 25 μL Q5 polymerase, and 0.5 μM of NP0492 and NP0546 primers. 2) During Ampure XP cleanup, samples were eluted into 21 μL water, and 40% (8 μL) was used in the second-round PCRs. 3) During the second-round PCRs, samples were amplified for a maximum of 20 cycles or until all samples were above 3,000 relative fluorescence units. 4) Samples were sequenced using custom sequencing primers NP0494, NP0495, NP0497, and NP0550 on either a NextSeq 550 using a NextSeq 500/550 High Output v2.5 75 cycle kit or on a NextSeq 2000 using a NextSeq 1000/2000 P3 50 cycle kit (Illumina).

### Calculating scores and classifications

Using a custom script, barcode sequencing reads were first converted to FASTQ format and demultiplexed using bcl2fastq (v2.20). Next, forward and reverse barcode reads were paired using PEAR (v0.9.11), and unique barcodes were counted^150^. To reduce error associated with counting noise, we removed low frequency variants below a threshold of 1 x 10^-6^ as well as variants that were observed in fewer than two replicates, which ultimately eliminated 19 variant effect scores (**Supplementary Table 1**). FIX variants were then assigned to sequenced barcodes using the barcode-variant map from PacBio sequencing. Any barcode associated with insertions, deletions, or multiple amino acid substitutions in FIX was removed from the analysis.

Secretion and γ-carboxylation scores were calculated by using a modification of a previous analysis pipeline^33,41,58^. Briefly, for each experiment, a weighted average of every variant’s frequency in each bin was calculated with the following weights: *w_bin 1_*: 0.25, *w_bin 2_*: 0.5, *w_bin 3_*: 0.75, *w_bin 4_*: 1. The weighted average for each variant was then min-max normalized such that the median score of WT barcodes was 1 and the median score of the lowest 5^th^ percentile of missense variants was 0. The final secretion and γ-carboxylation scores for each variant were then averaged across all replicates. Standard errors for each score were calculated by dividing the standard deviation of the min-max normalized values for each variant by the square root of the number of replicate experiments in which it was observed.

### Description of computational methods

VAMP-seq abundance scores for PTEN (urn:mavedb:00000013-a-1), TPMT (urn:mavedb:00000013-b-1), VKOR (urn:mavedb:00000078-b-1), CYP2C9 (urn:mavedb:00000095-b-1), and NUDT15 (urn:mavedb:00000055-a-1) were downloaded from MAVEdb^153^. Gla domain protein sequences for human coagulation factor IX (P00740), human prothrombin (coagulation factor II, P00734), human coagulation factor VII (P08709), human coagulation factor X (P00742), human protein C (P04070), human protein S (P07225), human osteocalcin (bone gla-protein, P02818), bovine osteocalcin (bone Gla-protein, P02820), human growth arrest-specific protein 6 (P14393), *Pseudechis prophyriacus* venom prothrombin activator porpharin-D (P58L93), *Notechis scutatis* venom prothrombin activator notecarin-D1 (P82807), and *Oxyuranus scutellatus* venom prothrombin activator oscutarin-C (P58L96) were downloaded from UniProt and aligned using MUSCLE^154^. Signal peptide predictions were generated by submitting variant and WT FIX sequences to the SignalP 6.0 server^50^. ConSurf predictions were generated by submitting WT FIX sequence to the ConSurf server^59^.

### Clinical variant curation

Our control set of pathogenic and benign FIX missense variants were collected from ClinVar (https://www.ncbi.nlm.nih.gov/clinvar/; accessed 1/10/2023), MLOF, and gnomAD v.4.1^1,3,20^ (https://gnomad.broadinstitute.org/; accessed 12/3/2024). MLOF variants have been previously deposited into the EAHAD FIX clinical database (https://dbs.eahad.org/FIX), the CDC CHBMP database (https://www.cdc.gov/hemophilia/mutation-project/index.html), and published^20,23^. A complete set of MLOF variants used in this study, along with relevant information about these variants, are provided in **Supplementary Table 4**. ClinVar variants with conflicting classifications or if evidence was not provided were removed. As the incidence of hemophilia B is approximately 3.8 to 5 per 100,000 male births, we deemed any gnomAD FIX variants with minor allele frequency in hemizygotes of greater than 1 per 1,000 as benign^33,92,155,156^. Lastly, *F9* variants found in individuals tested in MLOF were deemed benign if clinical testing demonstrated normal FIX activity in a hemizygous male. The final curated dataset contained 135 pathogenic or likely pathogenic variants and 14 benign or likely benign variants, sufficient for classification model training purposes^6,98,99^.

Data on FIX antigen levels, FIX activity, and disease severity reported in individuals with hemophilia B were collected from EAHAD (accessed 10/9/2023) and collapsed based on variant, resulting in 594 final variants with at least one of the three clinical data points^23^ (**Supplementary Table 9**). Because FIX antigen and activity are often reported as “<1%”, “<1 IU/dL”, or “undetectable” when below the limit of detection in clinical assays, we gave variants with these clinical assay results a value of 0.1% for the purposes of our analyses. Values for FIX antigen or FIX activity were averaged across reported individual values for each variant. For disease severity classifications, a consensus-based approach was used to determine final disease severity. If there was a tie between two or more severities, the variant was removed from the analysis. Further, if a variant’s antigen, activity, or severity was only reported in a single individual, that variant was removed from further analysis. For statistical analysis, a Kruskal-Wallis test followed by post-hoc Dunn’s test to detect differences in FIX antigen and MultiSTEP functional scores across severities was performed.

### Random forest classifier

We built a random forest classifier to distinguish between true benign and pathogenic variants using the ranger and tidymodels packages in R^157^. First, benign/likely benign and pathogenic/likely pathogenic classes were combined. We then split the curated variants so 75% of the data was used for training and 25% was used for testing the performance of the final model. To account for unbalanced class sizes in our training dataset (8.4% benign/likely benign and 90.6% pathogenic/likely pathogenic), we performed class-based random oversampling (ROSE) and trained the model using 5-fold cross-validation^158^. Model performance was evaluated using ROC curves generated with the reserved test set. The final model was applied to 103 VUS from MLOF that were observed in our experimental data. We used the ACMG rules-based guidelines to reinterpret variant classifications using *in vitro* functional data as strong evidence of pathogenicity^4,20^. As an alternative, we also employed a Bayesian framework to ACMG rules-based guidelines^6,98,99^. We calculated an odds of pathogenicity (OddsPath) and applied *in vitro* functional data as moderate evidence of pathogenicity. All other evidence codes were applied according to ACMG adapted for an X-linked inherited monogenic mendelian disorder^20^. All 103 VUS and their updated classifications are provided in **Supplementary Table 4**.

## Supporting information

Supplemental Tables (zipped)

## Data and code availability

*F9* variant scores are available in **Supplementary Table 10** and at MaveDB (www.https://www.mavedb.org/) with accession number “urn:mavedb:00001200”. Raw Illumina sequencing, PacBio sequencing, and barcode-variant maps, and variant scores are available in the NCBI Gene Expression Omnibus (GEO) repository with accession number GSE242805. All other source data and code to reproduce figures and analyses presented in this work are available on GitHub at https://github.com/fowlerlab/multistep.

## Acknowledgements

We thank A.P. Leith, C. Lee, D.E. Prunkard, and A. Silvestroni of the UW Foege Flow Lab and the UW Pathology Flow Cytometry Core for their assistance with cell analysis, staining, and sorting; K.M. Munson of the UW PacBio Sequencing Service for assistance with long-read PacBio sequencing; D.A. Nickerson, L.M. Starita, D.J. Maly, S. Nariya, J.J. Stephany, and A.E. McEwen for advice on analyzing data and feedback on the manuscript. We thank S.W. Pipe and A. Scheller of the University of Michigan Department of Pediatrics and Department of Hematology for providing FVIII constructs and advice on FVIII expression. We thank J. Kulman for discussions of FIX carboxylation. We thank R. Kruse-Jarres for her commitment to support research that improves the lives of people living with bleeding disorders. We thank and acknowledge B.A. Konkle the PI of MLOF, the MLOF partners at Bloodworks, the American Thrombosis and Hemostasis Network, the National Hemophilia Foundation (now the National Bleeding Disorders Foundation), funding from Biogen/Bioverativ, providers and staff at HTC sites, and the 11,341 participants who made MLOF a success. This work was supported by the National Heart, Lung, and Blood Institute (R01HL152066 to J.M.J. and D.M.F., F30HL151075 to N.A.P., and R01HL149855 to J.P.S.), the National Human Genome Research Institute (RM1HG010461 and UM1HG011969 to D.M.F.), the National Institute of General Medical Sciences (R01GM109110 to D.M.F.), and the Washington Center for Bleeding Disorders (to J.M.J).

## Author contributions

N.A.P., K.W.L., J.P.S., J.M.J., and D.M.F. conceived of the work. N.A.P., J.P.S., J.M.J., and D.M.F. wrote the manuscript. N.A.P., R.L.P., M.K.W, B.D.Z., K.J.H., K.M.S., X.W., K.W.L., and A.T.C. performed experiments. N.A.P. performed the statistical and computational analysis. N.A.P., S.N.F., and J.M.J. manually curated ClinVar, gnomAD, and MLOF for variants. N.A.P. and A.F.R. prepared results for distribution. N.A.P., S.N.F., S.F., and J.M.J. performed variant reinterpretation. All authors approved the final manuscript.

---

**Supplementary Figure 1:**
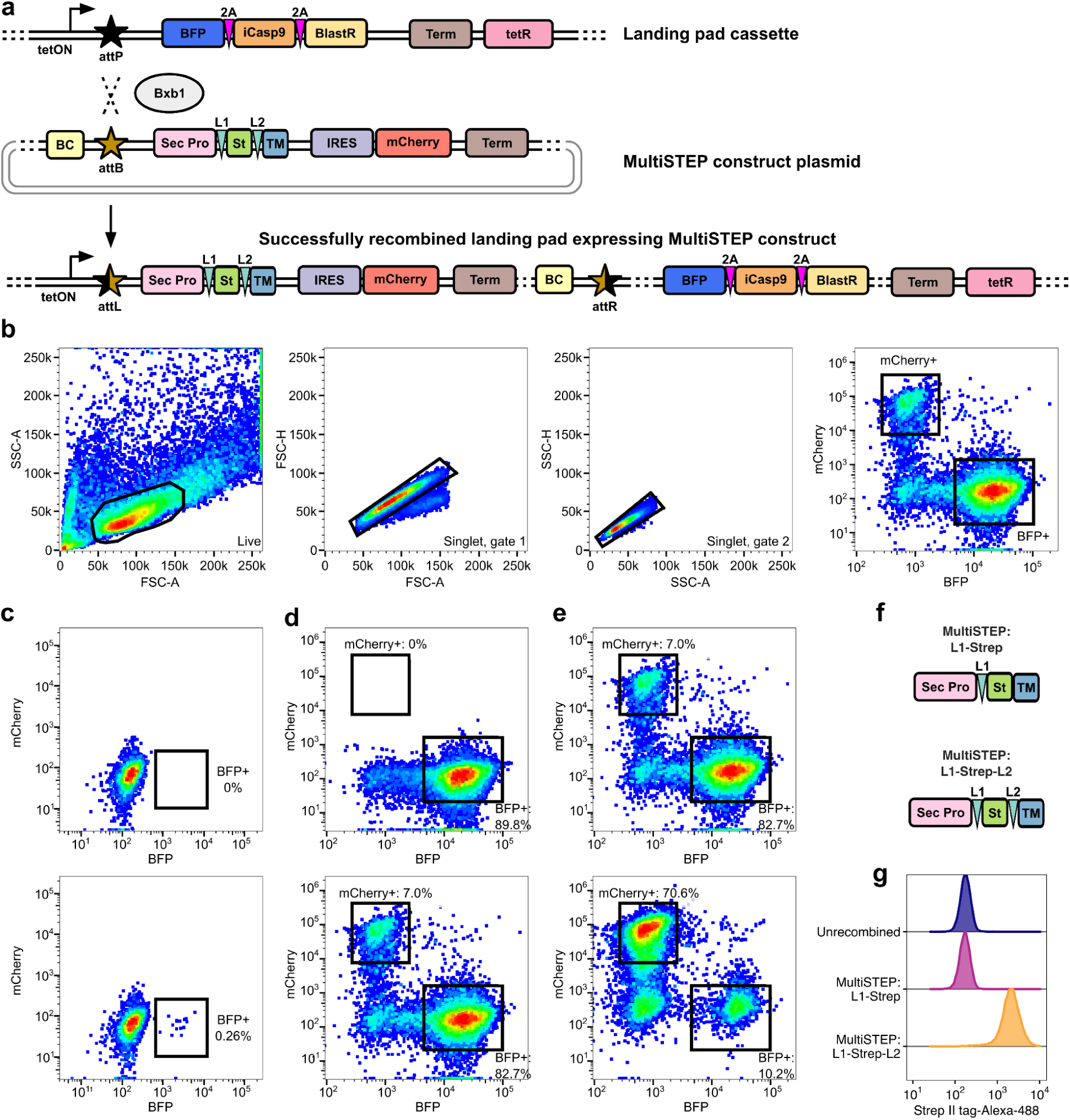
MultiSTEP is based on a flexible genomically integrated approach for expressing secreted protein variants. **a.** Cartoon depicting integration of a MultiSTEP plasmid construct into a genomically integrated landing pad cassette^31^. (Top): Lentivirally integrated landing pad cassette. Landing pad cassette expresses a BFP fluorophore (royal blue) from a tetON inducible promoter. The BFP fluorophore is fused to an inducible caspase-9 (iCasp9, orange) kill switch and a blasticidin resistance gene (dark yellow) with 2A sequences (neon pink) expressing BFP-2A-iCasp9-2A-BlastR from a tetON doxycycline-inducible promoter with a attP serine recombinase recognition site (black). Downstream is a terminator sequence (Term, brown) and tet repressor (tetR, salmon). Bxb1 serine recombinase, expressed from another plasmid, is shown in grey. (Middle): MultiSTEP plasmid construct. Secreted protein coding sequence (pink) is C-terminally fused to (GGGGS)2 flexible linkers (L1 and L2, teal) flanking a strep II epitope tag for surface detection (green) and a single pass CD28 transmembrane domain (TM, medium blue). An IRES (purple) drives co-transcription of an mCherry fluorophore (red) that serves as a transcriptional control. A downstream terminator sequence (Term, brown) is included. Upstream is an attB serine recombinase recognition sequence (goldenrod) and a unique 18 nucleotide degenerate barcode (BC, light yellow). (Bottom): Landing pad following plasmid integration. The attP and attB sequences have been recombined, forming attL and attR sequences (half-black, half-goldenrod). **b.** Sequential flow cytometry gating scheme for detecting and isolating landing pad cells that have integrated a MultiSTEP plasmid. Dot pseudocolor indicates density of cells. Sequential gates are outlined in black. BFP+, mCherry-cells (rightmost panel) represent unrecombined landing pad cells. mCherry+, BFP-cells represent successful recombinants. FSC: Forward scatter; SSC: side scatter. **c.** Comparison of 293-F cells with (bottom) and without (top) addition of lentivirus encoding the landing pad cassette using the gating scheme depicted in (**b**). Following lentivirus transduction, a polyclonal population of BFP+ 293-F cells were isolated via FACS and used for all experiments. Percentage of all cells within the indicated gate (n >10,000 cells overall). **d.** Comparison of polyclonal unrecombined landing pad 293-F cells (top) with cells transfected with a MultiSTEP plasmid encoding WT FIX (bottom) 7 days after transfection and 5 days after induction with doxycycline using the gating strategy outlined in (**b**). Number indicates the percentage of all cells within the adjacent gate (n > 10,000 cells). **e.** Comparison of cells transfected with a MultiSTEP construct encoding WT FIX 7 days after transfection and 5 days after induction with doxycycline cells that received only doxycycline (top) and cells that received doxycycline and 10 nM AP1903 (bottom). AP1903 activates the iCasp9 kill switch and enriches for successfully recombined cells. Number indicates the percentage of all cells within the adjacent gate (n >10,000 cells). Gating strategy as outlined in (**b**). **f.** Various MultiSTEP constructs. Secreted protein coding sequences (pink) are cloned into an attB-containing landing pad donor plasmid in (**a**). Secreted proteins are engineered to have C-terminally fused (GGGGS)_2_ flexible linkers (L1 and/or L2, teal) attached to a single pass transmembrane domain (TM, blue). In between the linkers is a strep II epitope tag for surface detection (green). MultiSTEP: L1-Strep does not contain an L2 linker, whereas MultiSTEP: L1-Strep-L2 does. **g.** Experimental flow cytometry of known well-secreted (A37T, S220T, WT) and poorly-secreted (C28Y) FIX variants displayed using MultiSTEP constructs depicted in (**f**) (n ∼30,000 cells per variant). Unrecombined cells do not display FIX and serve as a negative control. Fluorescent signal was generated by staining the library with a mouse monoclonal anti-strep II tag antibody followed by staining with an Alexa Fluor-647-labeled goat anti-mouse secondary antibody.

**Supplementary Figure 2:**
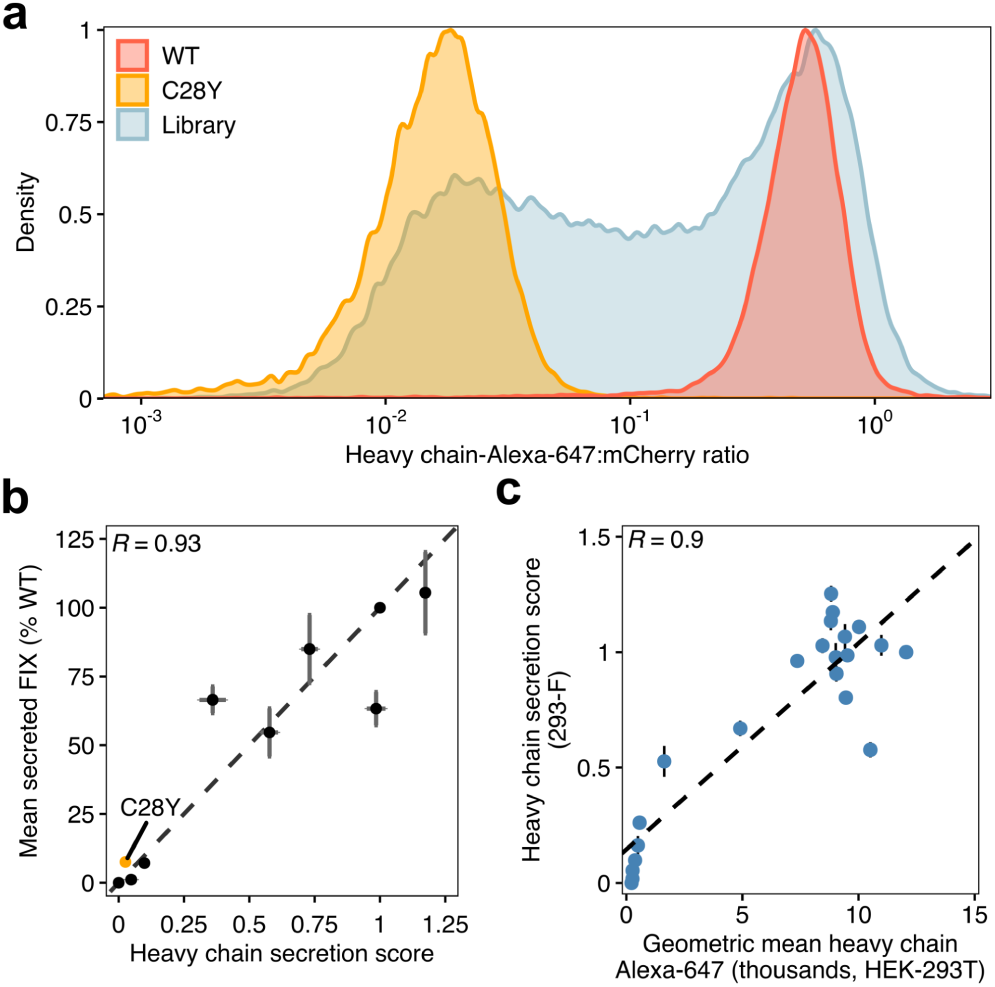
MultiSTEP-derived FIX secretion scores correlate with orthologous measures of FIX secretion. **a.** Flow cytometry of p.C28Y and WT controls (n = 10,000 cells) with the FIX library (n = 100,000 cells). p.C28Y overlaps the lowest mode of the library. **b.** Enzyme-linked immunosorbent assay quantification of secretion of eight untethered FIX missense variants (p.C28Y, p.A37T, p.G58E, p.E67K, p.G125V, p.C134R, p.S220T, and p.H267L) from 293-F cells compared to MultiSTEP-derived heavy chain secretion scores. p.C28Y shows negligible secretion from 293-F cells by both assays. Error bars show the standard error of the mean. Black dashed line indicates the line of perfect correlation between secretion scores. Pearson’s correlation coefficient is shown. **c**. Scatter plot comparing MultiSTEP-derived heavy chain secretion scores for 20 different FIX missense variants, WT, and unrecombined negative control to the geometric mean of Alexa Fluor-647 fluorescence measured using flow cytometry on cells expressing each variant individually. Error bars show standard error of the mean (n = 10,000 cells). Black dashed line indicates the line of best fit. Pearson’s correlation coefficient is shown.

**Supplementary Figure 3:**
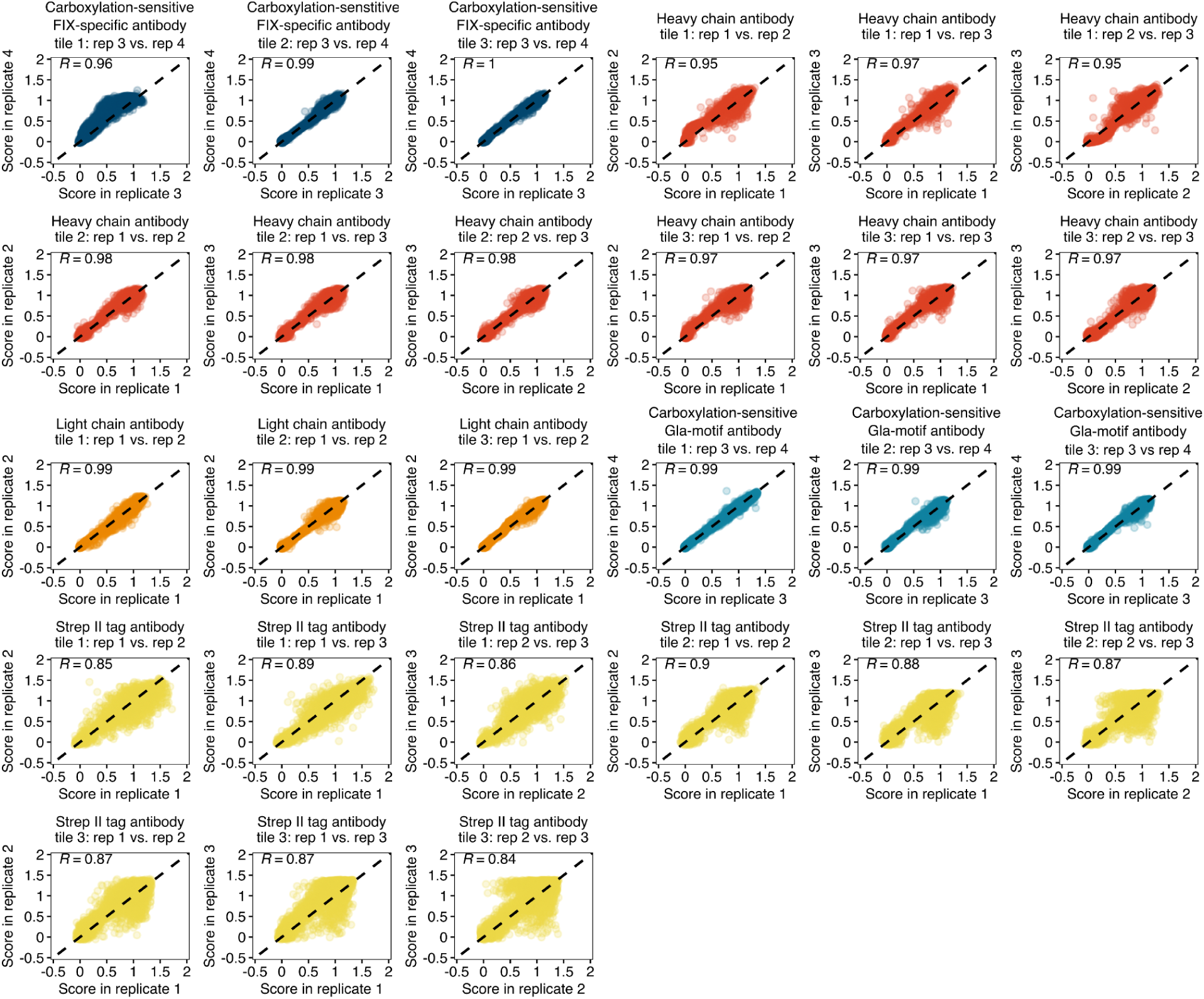
Correlations between FIX sublibrary tile experimental replicates for all antibodies. Pairwise MultiSTEP secretion and γ-carboxylation functional score correlations between replicate sorting experiments for FIX sublibrary tiles (n ≥ 3,144). Pearson’s correlation coefficients are shown.

**Supplementary Figure 4:**
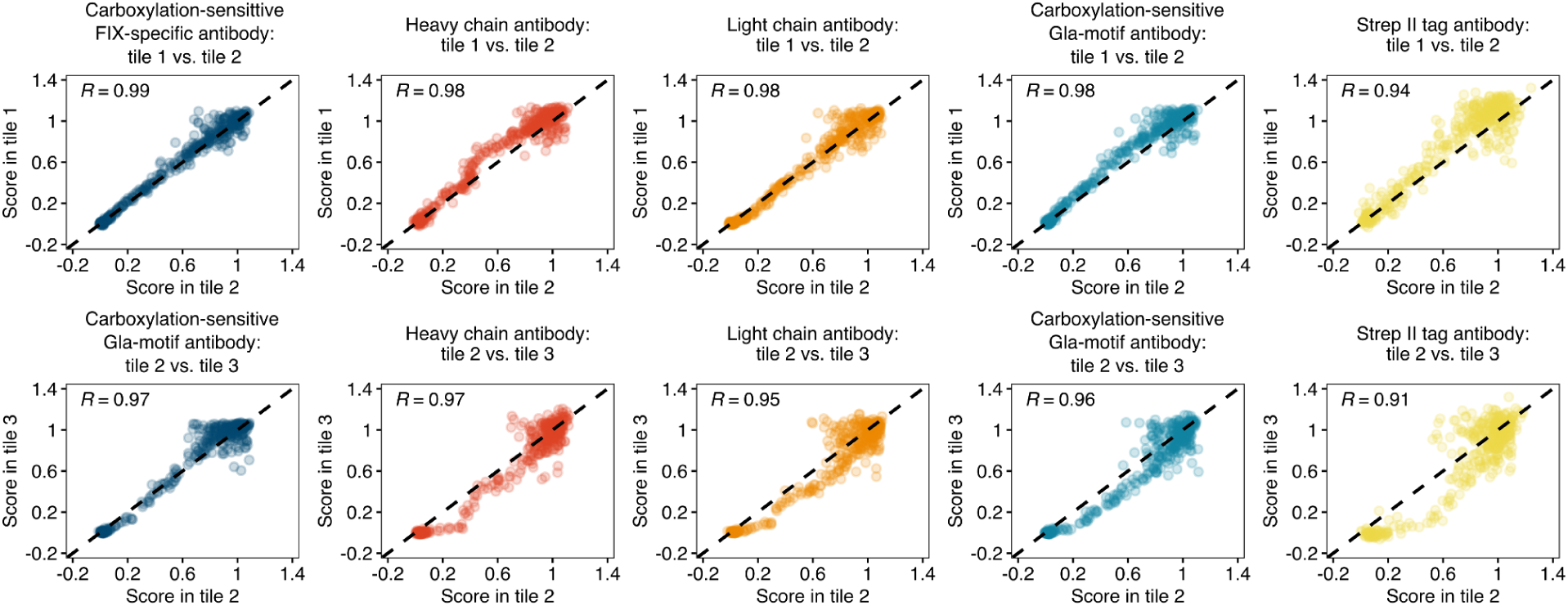
Correlations between FIX variants shared across multiple sublibrary tiles for all antibodies. Pairwise MultiSTEP secretion and γ-carboxylation functional score correlations for variants that are shared between sublibrary tiles (top row: n = 397, bottom row: n = 380). Pearson’s correlation coefficients are shown.

**Supplementary Figure 5:**
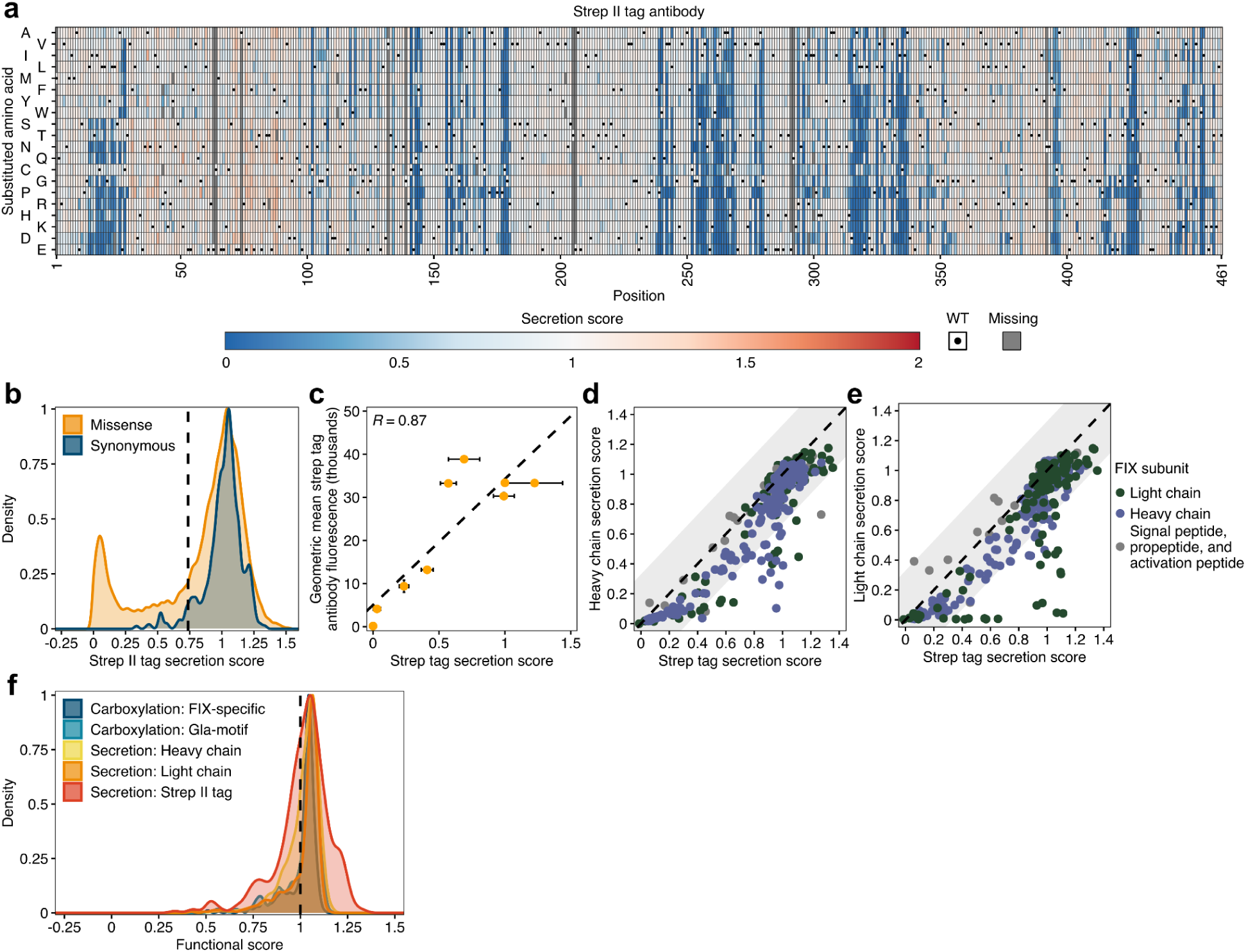
A flexible tag-based approach to assessing variant effects on secretion. **a.** Heatmap showing strep tag secretion scores for nearly all missense FIX variants. Heatmap color indicates antibody score from 0 (blue, lowest 5% of scores) to white (1, WT) to red (increased antibody scores). Black dots indicate the WT amino acid. Missing data scores are colored gray. **b.** Density distributions of strep tag secretion scores for FIX missense variants (orange) and synonymous variants (blue). Dashed line denotes the 5th percentile of the synonymous variant distribution. **c.** Scatter plot comparing MultiSTEP-derived strep tag secretion scores for seven different variants to the geometric mean of Alexa Fluor-647 fluorescence measured using flow cytometry on cells expressing each variant individually. Error bars show standard error of the mean (n = 3; >10,000 cells per replicate). Black dashed line indicates the line of best fit. Pearson’s correlation coefficient is shown. **d-e.** Scatter plots of median MultiSTEP-derived strep tag secretion scores vs. heavy chain (**d**) or light chain (**e**) at each position in FIX. Points are colored by chain architecture, using the same color scheme as Fig. 2a. Black dashed line indicates the line of perfect correlation between secretion scores. Gray background indicates <0.3 point deviation from perfect correlation. **f**. Density plots of MultiSTEP-derived synonymous variant scores generated with the indicated antibody. The dashed vertical line corresponds to a score of 1, equal to the WT score.

**Supplementary Figure 6:**
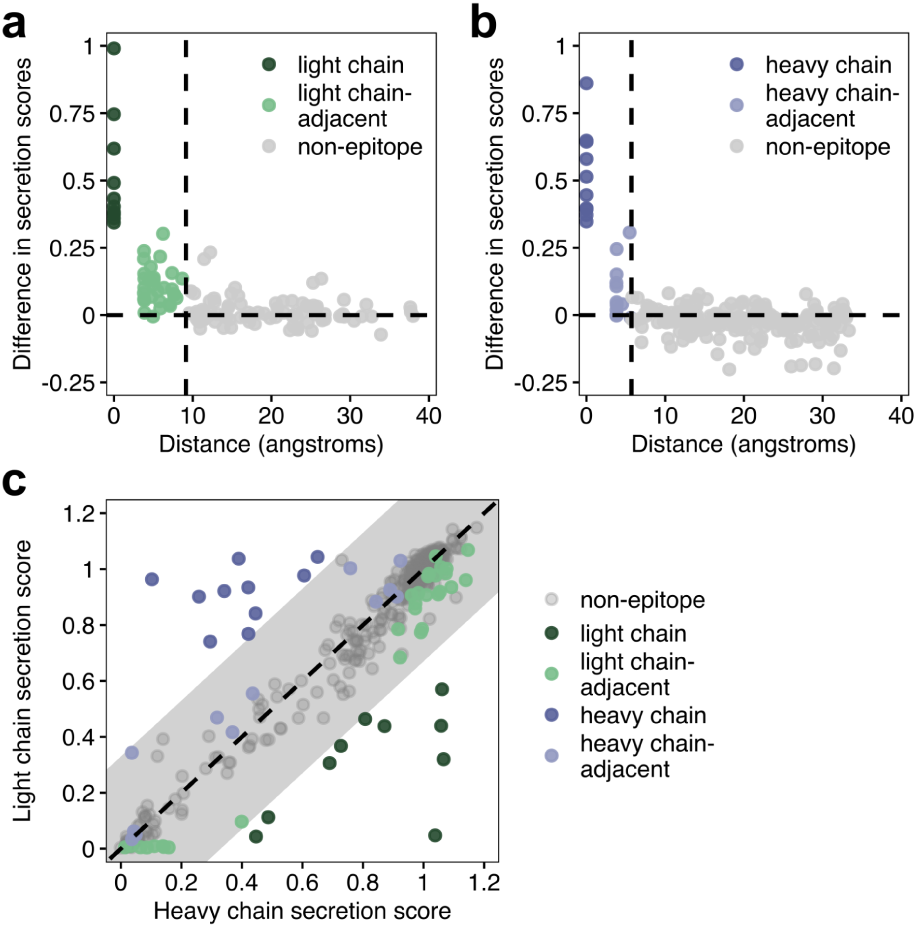
Variants near antibody epitopes demonstrate minor effects on secretion scores. **a.** Scatter plot of the difference in heavy chain and light chain secretion scores and the distance in angstroms between all α-carbons in the light chain and the nearest light chain epitope α-carbon using the AlphaFold2 model of mature, two-chain FIX. Low-confidence positions with predicted local distance difference test score (pLDDT) of <70 were removed from analysis. Color indicates whether a position was identified in the light chain epitope in Fig. 2h. Horizontal dashed line indicates a difference score of 0 (i.e., no difference in secretion scores). Vertical dashed line indicates boundary of likely epitope-adjacent effects on secretion scores as determined by changepoint analysis (9.15 angstroms). **b.** Scatter plot of the difference in heavy chain and light chain secretion scores and the distance in angstroms between all α-carbons in the heavy chain and the nearest heavy chain epitope α-carbon using the AlphaFold2 model of mature, two-chain FIX. Low-confidence positions with pLDDT of <70 were removed from analysis. Color indicates whether a position was identified in the heavy chain epitope in Fig. 2h. Horizontal dashed line indicates a difference score of 0 (i.e., no difference in secretion scores). Vertical dashed line indicates boundary of likely epitope-adjacent effects on secretion scores as determined by changepoint analysis (5.71 angstroms). **c.** Scatter plot of median MultiSTEP-derived heavy chain and light chain secretion scores at each position in FIX. Points are colored by whether they were identified as an epitope position (Fig. 2h) or an epitope-adjacent position as in (**a**) and (**b**). Black dashed line indicates the line of perfect correlation between secretion scores. Gray background indicates <0.3 point deviation from perfect correlation.

**Supplementary Figure 7:**
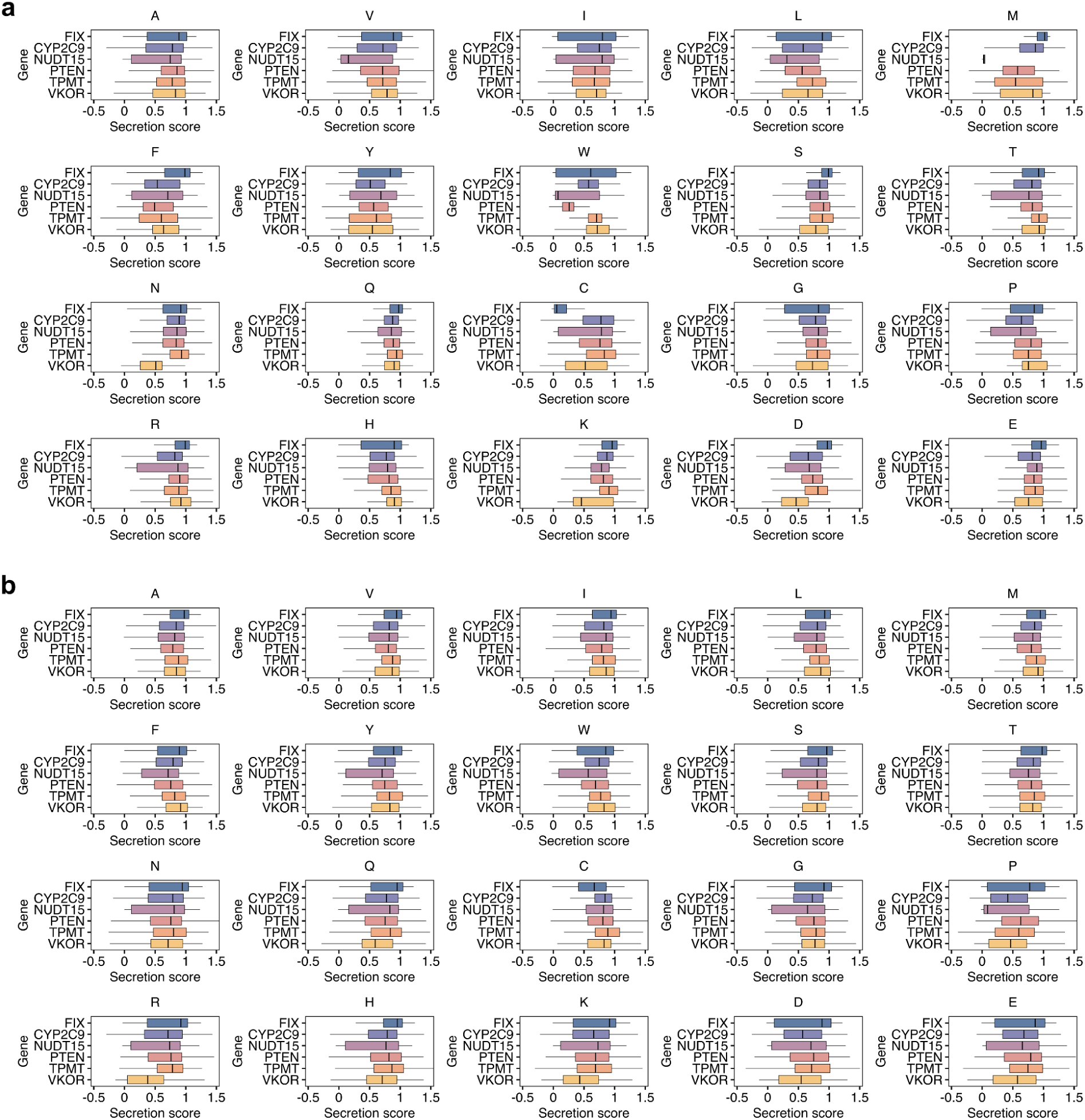
Effect of missense FIX variation on secretion compared to missense variant effects on abundance in cytosolic or transmembrane proteins. **a.** Box plots of secretion or abundance scores for all nonsynonymous variants across all positions with the indicated WT amino acid for six different proteins. For FIX, the average secretion score determined by MultiSTEP is shown. For all other proteins, the average abundance score measured using VAMP-seq is shown^33,40,41,58^. **b.** Box plots of secretion or abundance scores for all nonsynonymous substitutions of the indicated variant amino acid across all positions for six different proteins. Average scores were determined as in (**a**).

**Supplementary Figure 8:**
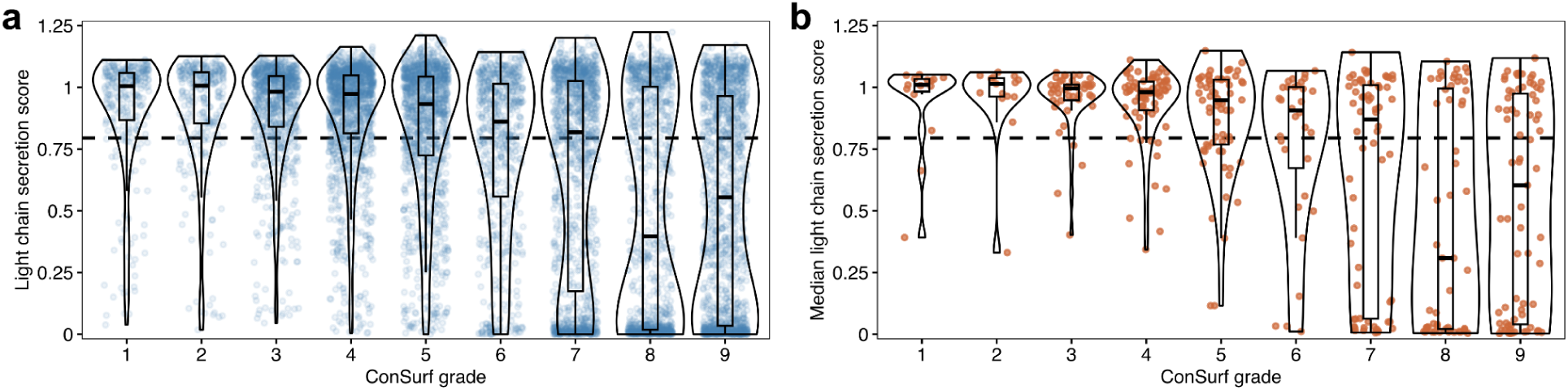
Sequence conservation strongly influences the effect of variation on FIX secretion. **a.** Comparison of light chain secretion scores with conservation grades from ConSurf^59^ (1: least conserved, 9: most conserved). Violin plot shows distribution of points with an inset box plot and horizontal lines representing the 25th, 50th, and 75th percentiles. Dashed horizontal line is the 5th percentile of the synonymous secretion score distribution. **b.** Comparison of median light chain secretion scores with conservation grades from ConSurf (1: least conserved, 9: most conserved). Violin plot shows distribution of points with an inset box plot and horizontal lines representing the 25th, 50th, and 75th percentiles. Dashed horizontal line is the 5th percentile of the synonymous secretion score distribution.

**Supplementary Figure 9:**
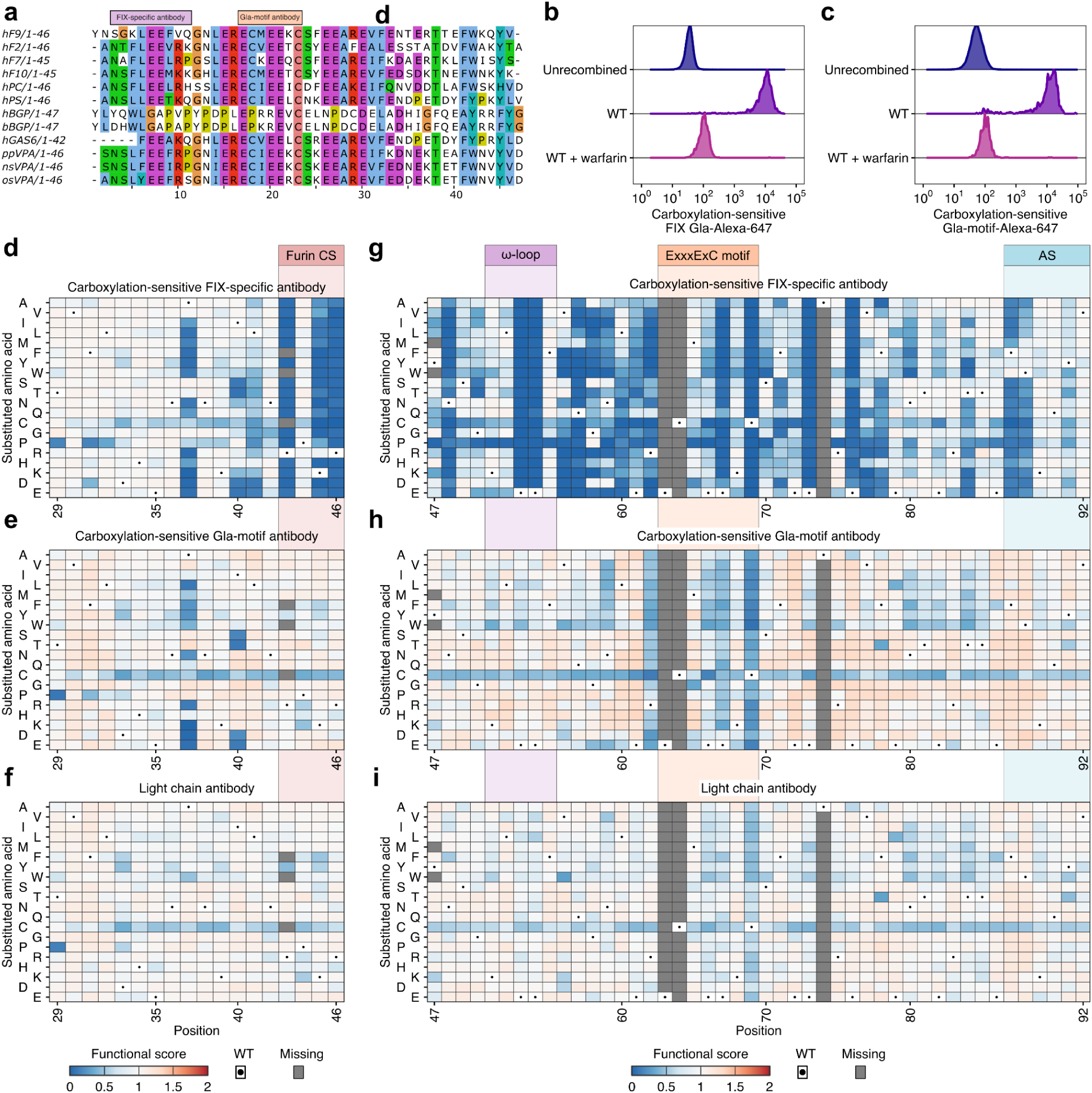
Carboxylation-sensitive antibodies identify functional motifs. **a.** Multiple sequence alignment of Gla-domain containing proteins that bind the carboxylation-sensitive Gla-motif (ExxxExC) antibody using MUSCLE^154^. Sequences were retrieved from UniProt. Antibody epitopes for both the carboxylation-sensitive FIX-specific antibody (ω-loop) and the carboxylation-sensitive Gla-motif antibody are shown. hF9: human coagulation factor IX (P00740); hF2: human prothrombin (coagulation factor II, P00734); hF7: human coagulation factor VII (P08709); hF10: human coagulation factor X (P00742); hPC: human protein C (P04070); hPS: human protein S (P07225); hBGP: human osteocalcin (bone gla-protein, P02818); bBGP: bovine osteocalcin (bone Gla-protein, P02820); hGAS6: human growth arrest-specific protein 6 (P14393); ppVPA: *Pseudechis prophyriacus* venom prothrombin activator porpharin-D (P58L93); nsVPA: *Notechis scutatis* venom prothrombin activator notecarin-D1 (P82807); osVPA: *Oxyuranus scutellatus* venom prothrombin activator oscutarin-C (P58L96). **b-c.** Flow cytometry of WT FIX with and without pretreatment with warfarin in the presence of 50 nM vitamin K (n ∼30,000 cells per variant). Warfarin inhibits the activity of vitamin K epoxide reductase complex 1 (VKORC1), an enzyme essential in the steps leading to γ-carboxylation of FIX. Unrecombined cells do not display FIX and serve as a negative control. Fluorescent signal was generated by staining the library with either a mouse monoclonal carboxylation-sensitive FIX-specific antibody (**b**) or a mouse monoclonal carboxylation-sensitive Gla-motif antibody that binds the conserved motif in (**a**), followed by staining with an Alexa Fluor-647-labeled goat anti-mouse secondary antibody. **d-f.** Heatmaps showing carboxylation-sensitive FIX-specific carboxylation scores (**d**), carboxylation-sensitive Gla-motif carboxylation scores (**e**), or light chain secretion scores (**f**) for nearly all missense FIX variants in the propeptide. Furin cleavage site (Furin CS), ω-loop, ExxxExC motif, and aromatic stack (AS) are annotated above (**d**). Heatmap color indicates antibody score from 0 (blue, lowest 5% of scores) to white (1, WT) to red (increased antibody scores). Black dots indicate the WT amino acid. Missing data scores are colored gray. **g-i.** Heatmaps showing carboxylation-sensitive FIX-specific carboxylation scores (**g**), carboxylation-sensitive Gla-motif carboxylation scores (**h**), or light chain secretion scores (**i**) for nearly all missense FIX variants in the Gla domain. ω-loop, ExxxExC motif, and aromatic stack (AS) are annotated above (**g**). Heatmap color indicates antibody score from 0 (blue, lowest 5% of scores) to white (1, WT) to red (increased antibody scores). Black dots indicate the WT amino acid. Missing data scores are colored gray.

**Supplementary Figure 10:**
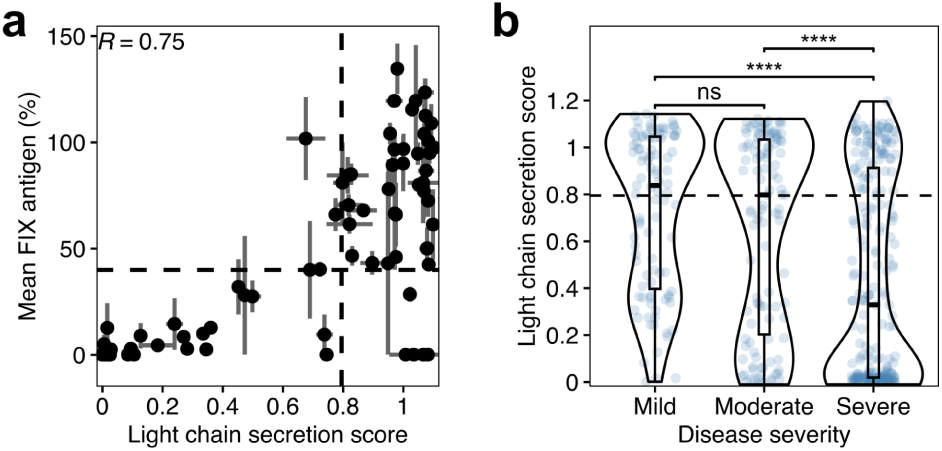
Removal of epitope-adjacent positions from analysis does not change key secretion score results. **a.** Scatter plot of light chain secretion scores and FIX plasma antigen from individuals with hemophilia B in the EAHAD database with light chain epitope-adjacent positions identified in **Supplementary Fig. 6a** removed (n = 19 variants). Horizontal solid lines indicate standard error of the mean for light chain secretion scores. Vertical solid lines indicate standard error of the mean for FIX plasma antigen levels across individuals with hemophilia B harboring the same variant. Dashed horizontal line is 40% FIX plasma antigen. Dashed vertical line is the 5th percentile of the synonymous secretion score distribution. **b.** Comparison of EAHAD individual hemophilia B severity with light chain secretion scores from MultiSTEP with light chain epitope-adjacent positions identified in **Supplementary Fig. 6a** removed (n = 40 variants). Violin plot shows distribution of points with an inset box plot representing the 25th, 50th, and 75th percentiles. Dashed horizontal line is the 5th percentile of the synonymous secretion score distribution. To compare median secretion scores across disease severities, a Kruskal–Wallis test adjusted for multiple comparisons by post-hoc Dunn’s test was performed. Asterisks indicate the level of statistical significance. ns = p >0.05. **** = p <0.0001.

**Supplementary Figure 11:**
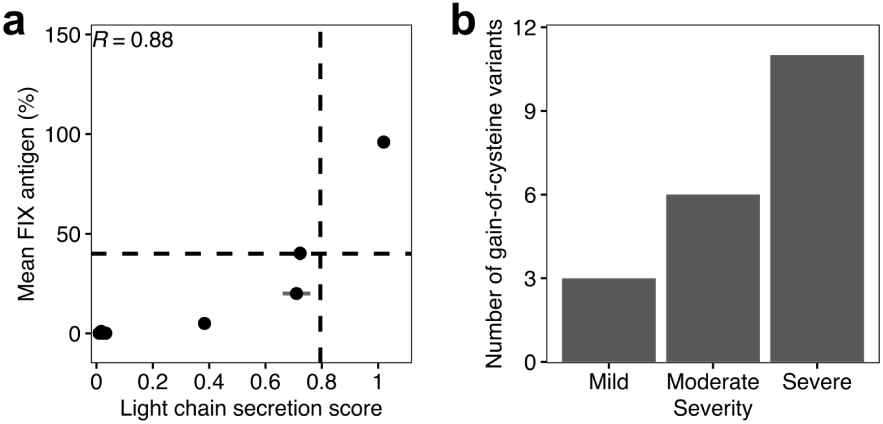
Clinical consequences of gain-of-cysteine variants. **a.** Scatter plot of light chain secretion scores and FIX plasma antigen from individuals with hemophilia B harboring gain-of-cysteine variants in the EAHAD database (n = 9). Horizontal solid lines indicate standard error of the mean for light chain secretion scores. Vertical solid lines indicate standard error of the mean for FIX plasma antigen levels across individuals with hemophilia B harboring the same variant. Dashed horizontal line is 40% FIX plasma antigen. Dashed vertical line is the 5th percentile of the synonymous secretion score distribution. **b.** Bar plot of hemophilia B disease severity in the EAHAD database for individuals harboring gain-of-cysteine variants.

**Supplementary Figure 12:**
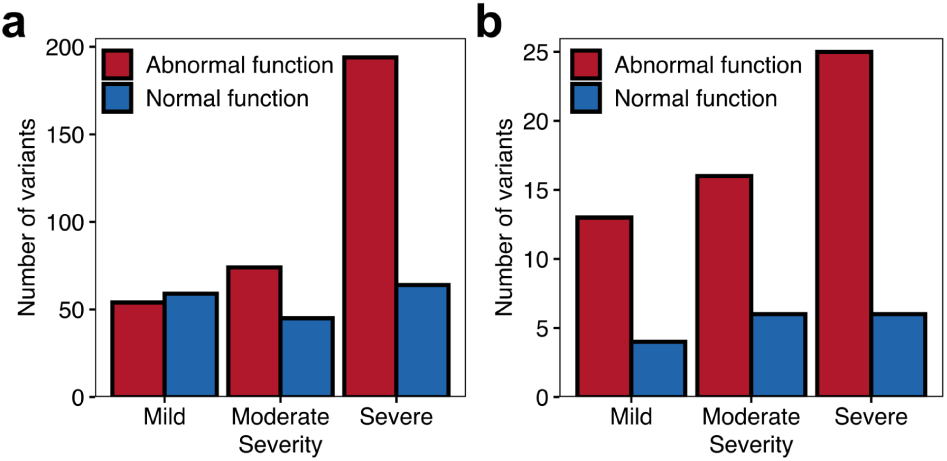
Random forest model predictions for FIX variants in the EAHAD FIX Variant Database associated with hemophilia B. **a.** Bar plot of the number of FIX variants in the EAHAD database and their classification using the random forest model trained on MultiSTEP functional data, by disease severity. Color indicates model prediction. **b.** Bar plot of the number of FIX propeptide and Gla domain variants in the EAHAD database and their classification using the random forest model trained on MultiSTEP functional data, by disease severity. Color indicates model prediction.

**Supplementary Figure 13:**
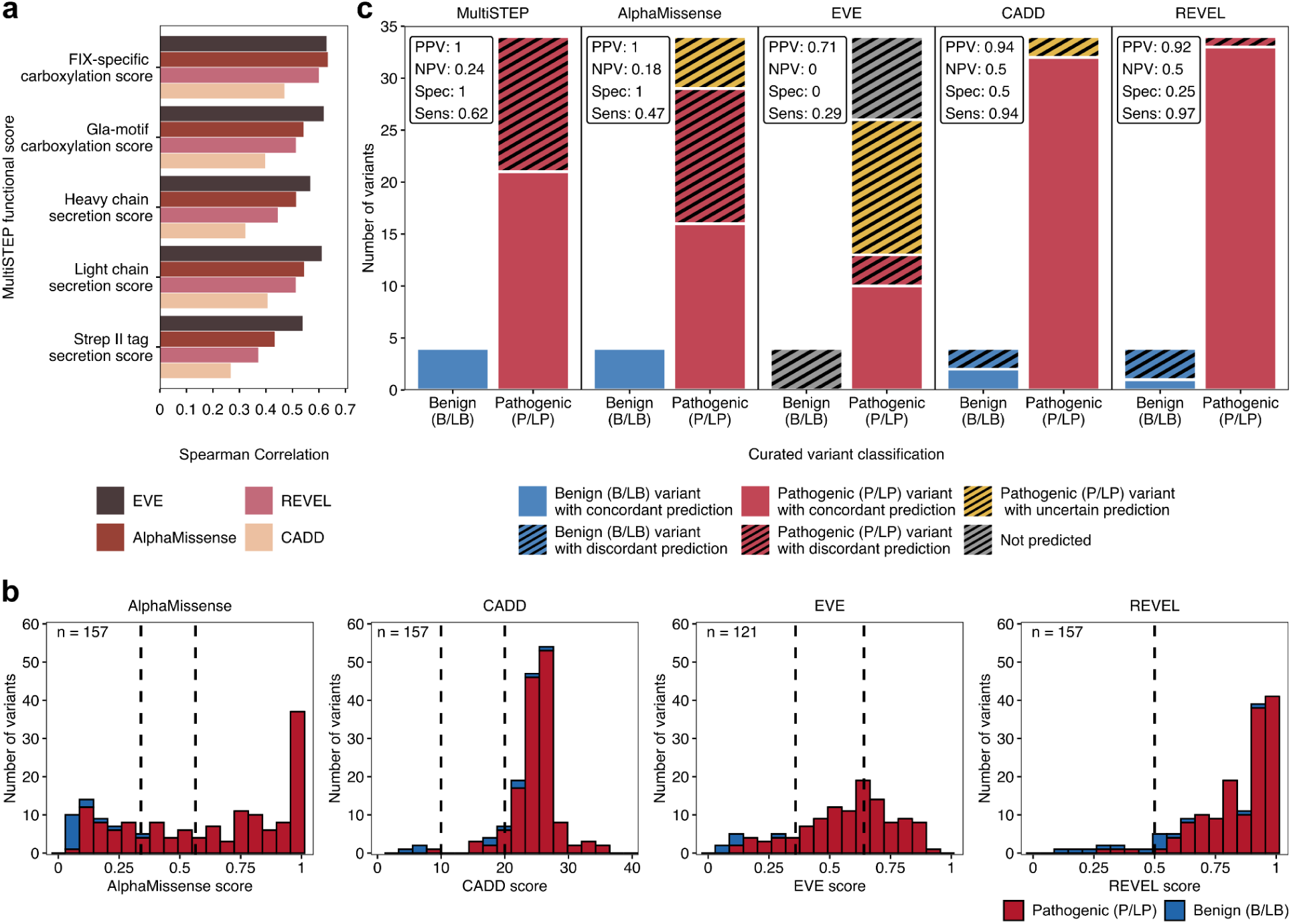
Random forest model predictions for FIX variants in the EAHAD FIX Variant Database associated with hemophilia B. **a.** Spearman correlation of MultiSTEP functional scores with EVE, AlphaMissense, REVEL, and CADD variant effect predictors. **b.** Histograms of four variant effect predictor scores for F9 missense variants of known effect curated from ClinVar, gnomAD, and MLOF. Color indicates clinical variant interpretation. Data from four variant effect predictors are shown. Black dashed vertical lines indicate the thresholds for each predictor. For AlphaMissense we used the thresholds recommended in the original publication for 90% precision on existing ClinVar annotated variants (≤0.34: benign, 0.34-0.564: uncertain, ≥0.564: pathogenic). For REVEL, we used the thresholds used in the initial publication to assess REVEL’s precision in ClinVar (<0.5: benign, 0.5: uncertain, >0.5 pathogenic). For EVE, we used the thresholds recommended in the original publication for the 75% most confident classifications (≤0.359: benign, 0.359-0.641: uncertain, ≥0.641: pathogenic). For CADD, we used the same thresholds used in the MLOF clinical laboratory (<10: benign, 10-20: uncertain, >20: pathogenic). Number of variants scored by each predictor is annotated. **c.** Classification accuracy for *F9* missense variants of known effect curated from ClinVar, gnomAD, and MLOF in the test set of variants (benign/likely benign, n = 4; pathogenic/likely pathogenic, n = 34) by MultiSTEP random forest model and four variant effector predictors using thresholds defined in (**b**). True benign/likely benign and pathogenic/likely pathogenic labels are denoted on the x-axis, and columns are colored relative to the classification for each method. Solid colors indicate correct classification, whereas striped colors indicate incorrect classification. For variant effect predictors, missing variants are colored gray with stripes and uncertain predictions are colored yellow with stripes. PPV: positive predictive value; NPV: negative predictive value; Spec: specificity; Sens: sensitivity.

**Supplementary Figure 14.**
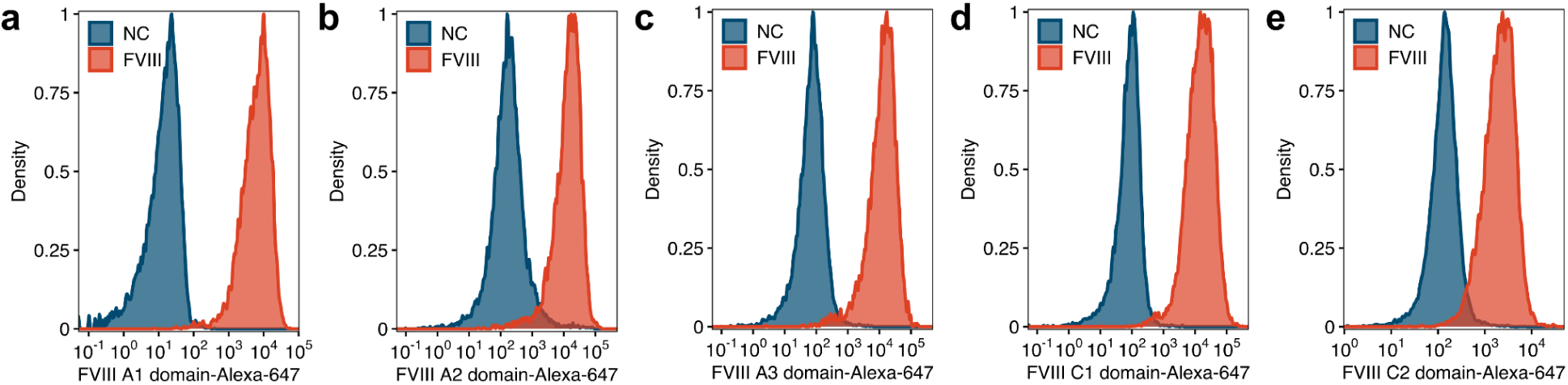
Detection of cell-surface displayed FVIII. Experimental flow cytometry of B-domain deleted coagulation factor VIII (FVIII) in the MultiSTEP backbone (n = ∼30,000 cells per variant). Unrecombined cells (NC) do not display FVIII and serve as a negative control. Fluorescent signal was generated by staining cells with mouse monoclonal anti-FVIII antibodies specific to each of the five FVIII domains in the heavy chain [A1 (**a**) and A2 (**b**)] and light chain [A3 (**c**), C1 (**d**), and C2 (**e**)], followed by staining with an Alexa Fluor-647-labeled goat anti-mouse secondary antibody.

**Supplementary Table 1: Library and assay statistics.**

Number of FIX variants assessed at various stages of library preparation and in each MultiSTEP assay.

**Supplementary Table 2: Variants with discordant circulating antigen and secretion scores.**

Variants with discordant secretion scores and FIX plasma antigen from individuals with hemophilia B in the EAHAD database^23^. Variants with undetectable FIX antigen are labeled as <1%, as reported by the clinical laboratory in EAHAD. Variants are classified as low antigen if they have a mean circulating FIX antigen of <40%. Variants are classified as low secretion if they have a mean secretion score that is less than 0.795, which is the 5th percentile of the synonymous secretion score distribution. SE: standard error of the mean.

**Supplementary Table 3: Random forest model classifications for 8,964 F9 variants.** Mean secretion scores, γ-carboxylation scores, and functional variant classifications made using the random forest classifier we trained on known pathogenic/likely pathogenic and benign/likely benign *F9* missense variants (see Methods). Variants without functional scores for all antibodies were removed prior to classifier implementation and do not have associated functional predictions.

**Supplementary Table 4: Classification criteria and reclassification results for My Life, Our Future *F9* variants.**

Classification criteria, random forest classifier predictions, and resultant pathogenicity classifications for 214 *F9* variants from *My Life, Our Future*^20^. Classification criteria (columns BP1 through PVS1 as defined by the American College of Medical Genetics and Genomics^4^) were used in variant curation by clinical experts in hemophilia genetics based on available clinical data, databases, and literature review. PVS1: Null variant (nonsense, frameshift, canonical ±1 or 2 splice sites, initiation codon, single or multi-exon deletion) in a gene where loss of function is a known mechanism of disease. PS1: Same amino acid change as a previously established pathogenic variant regardless of nucleotide change. PS2: *De novo* (both maternity and paternity confirmed) in a patient with the disease and no family history. PS3: Well-established *in vitro* or *in vivo* functional studies supportive of a damaging effect on the gene or gene product. PS4: The prevalence of the variant in affected individuals is significantly increased compared with the prevalence in controls. PM1: Located in a mutational hot spot and/or critical and well-established functional domain (e.g., active site of an enzyme) without benign variation. PM2: Absent from controls (or at extremely low frequency if recessive) in population databases. PM3: For recessive disorders, detected in trans with a pathogenic variant. PM4: Protein length changes as a result of in-frame deletions/insertions in a non-repeat region or stop-loss variants. PM5: Novel missense change at an amino acid residue where a different missense change determined to be pathogenic has been seen before. PM6: Assumed *de novo*, but without confirmation of paternity and maternity. PP1: Cosegregation with disease in multiple affected family members in a gene definitively known to cause the disease. PP2: Missense variant in a gene that has a low rate of benign missense variation and in which missense variants are a common mechanism of disease. PP3: Multiple lines of computational evidence support a deleterious effect on the gene or gene product (conservation, evolutionary, splicing impact, etc.). PP4: Patient’s phenotype or family history is highly specific for a disease with a single genetic etiology. PP5: Reputable source recently reports variant as pathogenic, but the evidence is not available to the laboratory to perform an independent evaluation. BA1: Allele frequency is >5% in population databases. BS1: Allele frequency is greater than expected for disorder. BS2: Observed in a healthy adult individual for a recessive (homozygous), dominant (heterozygous), or X-linked (hemizygous) disorder, with full penetrance expected at an early age. BS3: Well-established *in vitro* or *in vivo* functional studies show no damaging effect on protein function or splicing. BS4: Lack of segregation in affected members of a family. BP1: Missense variant in a gene for which primarily truncating variants are known to cause disease. BP2: Observed in trans with a pathogenic variant for a fully penetrant dominant gene/disorder or observed in cis with a pathogenic variant in any inheritance pattern. BP3: In-frame deletions/insertions in a repetitive region without a known function. BP4: Multiple lines of computational evidence suggest no impact on gene or gene product (conservation, evolutionary, splicing impact, etc.). BP5: Variant found in a case with an alternate molecular basis for disease. BP6: Reputable source recently reports variant as benign, but the evidence is not available to the laboratory to perform an independent evaluation. BP7: A synonymous (silent) variant for which splicing prediction algorithms predict no impact to the splice consensus sequence nor the creation of a new splice site and the nucleotide is not highly conserved. Original interpretations as well as reinterpretations using random forest classifier predictions as either moderate or strong evidence are included.

**Supplementary Table 5: Oligonucleotides.**

Oligonucleotides used in this study.

**Supplementary Table 6: FIX variant nomenclature.**

HGVS, legacy, and chymotrypsin numbering systems for each position in WT FIX. In chymotrypsin numbering, some position values in flexible loops that are not conserved repeat.

**Supplementary Table 7: Detailed antibody information.**

Properties, stock concentrations, and experimental conditions for each assayed antibody. Epitopes, when known, are provided.

**Supplementary Table 8: PCR replicate correlations.**

PCR replicate correlations for each genomic DNA-derived barcode library amplification.

**Supplementary Table 9: Case data for FIX variants in EAHAD.**

Variant-level data for FIX variants curated from the EAHAD FIX variant database (accessed 10/9/2023).

**Supplementary Table 10: MultiSTEP variant scores.**

Table of FIX secretion and γ-carboxylation scores for nearly all possible missense and synonymous FIX variants. Scores are derived from each of the five antibodies presented in this work. The number of unique variants scored for each antibody ranges between 8,961 and 8,964 (Strep II tag antibody: 8,961; Heavy chain antibody: 8,963; Light chain antibody: 8,964; Carboxylation-sensitive FIX-Gla antibody: 8,964; Carboxylation-sensitive Gla-motif antibody: 8,964). There are a total of 44,816 measured secretion and γ-carboxylation variant effects.

